# Recessive deleterious variation has a limited impact on signals of adaptive introgression in human populations

**DOI:** 10.1101/2020.01.13.905174

**Authors:** Xinjun Zhang, Bernard Kim, Kirk E. Lohmueller, Emilia Huerta-Sánchez

## Abstract

Admixture with archaic hominins has altered the landscape of genomic variation in modern human populations. Several gene regions have been previously identified as candidates of adaptive introgression (AI) that facilitated human adaptation to specific environments. However, simulation-based studies have suggested that population genetics processes other than adaptive mutations, such as heterosis from recessive deleterious variants private to populations before admixture, can also lead to patterns in genomic data that resemble adaptive introgression. The extent to which the presence of deleterious variants affect the false-positive rate and the power of current methods to detect AI has not been fully assessed. Here, we used extensive simulations to show that recessive deleterious mutations can increase the false positive rates of tests for AI compared to models without deleterious variants. We further examined candidates of AI in modern humans identified from previous studies and show that, although deleterious variants may hinder the performance of AI detection in modern humans, most signals remained robust when deleterious variants are included in the null model. While deleterious variants may have a limited impact on detecting signals of adaptive introgression in humans, we found that at least two AI candidate genes, *HYAL2* and *HLA*, are particularly susceptible to high false positive rates due to the recessive deleterious mutations. By quantifying parameters that affect heterosis, we show that the high false positives are largely attributed to the high exon densities together with low recombination rates in the genomic regions, which can further be exaggerated by the population growth in recent human evolution. Although the combination of such parameters is rare in the human genome, caution is still warranted in other species with different genomic composition and demographic histories.

## Introduction

Gene flow between populations can rapidly increase the genetic variation in the recipient group by introducing new variants from a different population. If some of this genetic variation increases an organism’s survival and reproduction, it can be considered adaptive. Adaptive introgression has been found to facilitate adaptation to local environments in a wide range of taxa, from plants to animals^1–4^. In modern humans, introgression with archaic hominins, including Neanderthals^5,6^ and Denisovans^7,8^, has changed the genomic diversity of and supplied adaptive alleles to most populations outside of Africa. Previous studies have identified at least 30 candidate genomic regions in modern humans that were putatively adaptively introgressed^9–19^ – among which one of the most well-known example is a Denisovan-like haplotype at the *EPAS1* gene that facilitated adaptation to high altitude in the Tibetan population^20,21^. As of today, the putative AI tracts in modern humans can be traced back to Neanderthals^9,18,19,22^, Denisovans^13,20^, unknown archaic groups^23,24^, or a mix of more than one population^1,22^.

The detection of adaptive introgression mostly relies on independently looking for signatures of introgression^22,24–27^ and signatures of positive selection^28–33^. Additionally, a number of allele frequency-based summary statistics have been shown to be particularly powerful at directly inferring AI without needing to apply separate tests for introgression or selection at genomic regions. These statistics include: the number of uniquely shared alleles between donor and recipient population (U statistic), the quantile distribution of derived alleles in recipient (Q statistic), and sequence divergence ratio (RD)^11^. Racimo *et al.*^11^ further demonstrated the robustness of these statistics to several factors that may confound the detection of AI, including incomplete lineage sorting and ancestral population structure.

While there is tremendous interest in identifying candidate regions for AI, most mutations that occur in genomes are likely either neutral or deleterious^34^. Deleterious mutations continue to accumulate in the distinct populations after they split from each other^35^. These deleterious mutations can also affect the genomic landscape in the recipient population after introgression. The genetic load (i.e. reduction in population fitness due to deleterious variants) of archaic hominins is usually higher than modern humans due to the former’s small effective population size^5^. Thus, most introgressed archaic ancestry is ultimately purged from the modern human gene pool^36,37^. Conversely, a higher frequency of archaic variants and longer introgressed tracts are the typical signatures indicating adaptive introgression. However, recent studies suggest that other population genetics processes can also generate long introgressed tracts at high frequencies in a recipient population. For example, if many deleterious mutations are recessive, and are private to one population^38–40^, after introgression homozygous recessive alleles (from either donor or recipient) will most likely become heterozygotes. In this situation, new haplotypes get created in the admixed population where the negative fitness effects on such variants are now reduced or eliminated. As such, an initial heterosis effect occurs (Fig. 1), since admixed individuals have higher fitness compared to unadmixed individuals due to the masking of recessive deleterious variants. The neutral markers nearby the recessive deleterious variants would also increase in frequency^41,42^, leading to an overall increase of introgressed ancestry in the admixed population^37^, resembling what is expected from adaptive introgression^1,11^.

**Figure 1:**
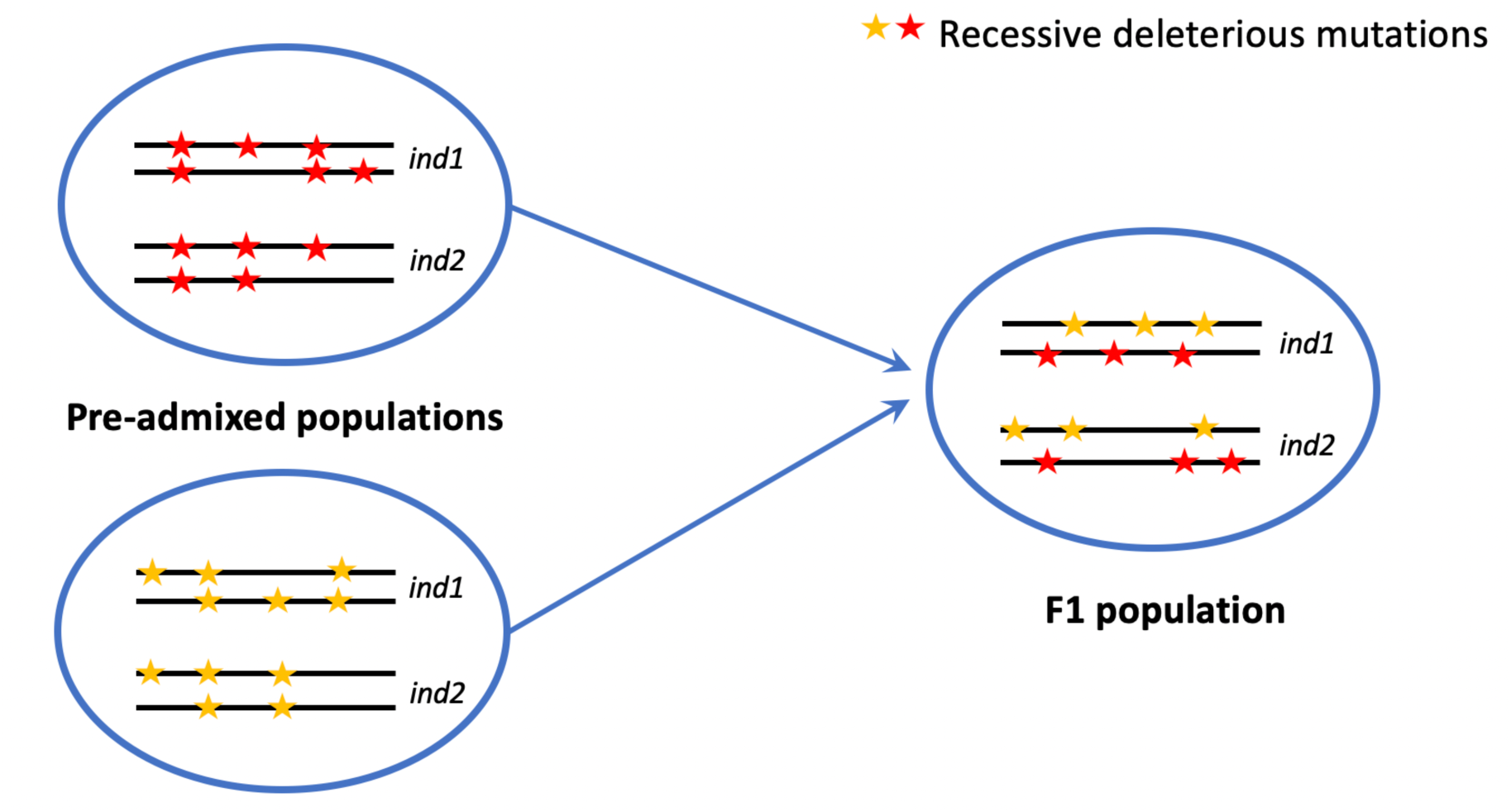
Heterosis effect from an increase in heterozygosity due to admixture. A red or yellow star represents a mutation that is deleterious and recessive (h=0). In this figure, each individual in the pre-admixed populations is homozygous for recessive deleterious variants at 2 distinct sites. In the F1 population, if the two populations admix in equally, all mutations that were private to the original populations and were previously homozygous are now heterozygous in the F1 population.

As an example of this, Harris and Nielsen^37^ simulated modern human-Neanderthal admixture, and suggested that the heterosis effect from recessive deleterious variants can increase the Neanderthal ancestry in modern humans by up to 3%. Kim *et al.*^43^ showed that low recombination rate, high exon densities, and small recipient population size can all amplify the effect of deleterious variants leading to an increase in introgressed ancestry. However, both Harris and Nielsen and Kim *et al.* illustrated the confounding effect of deleterious variants on adaptive introgression by directly tracking the introgressed ancestry from simulations. Although straightforward and convenient in simulation studies, introgressed ancestry is difficult to precisely measure with the empirical data. Thus, it remains unclear whether other summary statistics aimed to detect adaptive introgression are affected by the presence of deleterious variants.

Our present work aims to systematically explore the behavior of the summary statistics for detecting adaptive introgression in the presence of deleterious, recessive variants in realistic human demographic models. By performing extensive simulations under different evolutionary parameters (demography, recombination rate, and genic structure), we show that accounting for recessive deleterious mutations in the null model leads to an increase in false positive rates in most statistics due to the heterosis effect, with some statistics being more robust than others.

By examining the currently known AI candidate regions in modern humans, we show that at least several candidate genes previously identified as being under AI (*HYAL2*^14^ and *HLA* gene cluster^15^) may alternatively be false-positives due to the presence of deleterious variants. However, we also show that most of the human AI candidate genes cannot be explained by deleterious variants, suggesting they may be genuine targets of AI. We further show that in *HYAL2*^14^ and *HLA*, a combination of high exon density and low recombination rate is the main factor contributing to the high false positive rates in the two genes. The evolutionary history of humans, especially the recent rapid population growth, slightly increases the false positive rate as well. Despite the overall limited impact from recessive deleterious variants on AI signals in human populations, deleterious mutations remain a confounding factor for reliable AI detection in other organisms with certain combination of evolutionary parameters such as high exon density and low recombination rate. As such, effects from deleterious variants are not negligible and should be included in the null models for identifying candidate regions of AI.

## Results

### Simulations and measurements of adaptive introgression

We used the program SLiM 3.2.0^44^ to simulate different models of admixture. Each of the models consists of three populations: an ancestral population at equilibrium that splits into two sub-populations (pD for “donor population” and pO for “outgroup”), and one of the subpopulations subsequently splits again after a period of time (pO, and pR for “recipient population”). After the second split, a pulse of admixture occurred at 10% from pD uni-directionally into pR, lasting for one generation. Fig. 2 shows an illustration of the two demographic models used herein: 1) Model_0 (Fig. 2a) represents a demography where the recipient population size is 10 times smaller than the donor population size throughout the simulation; and 2) Model_h (Fig. 2b) represents an estimated demography for modern humans, with a single pulse of archaic admixture introduced to the non-African population^5–7,45,46^.

**Figure 2:**
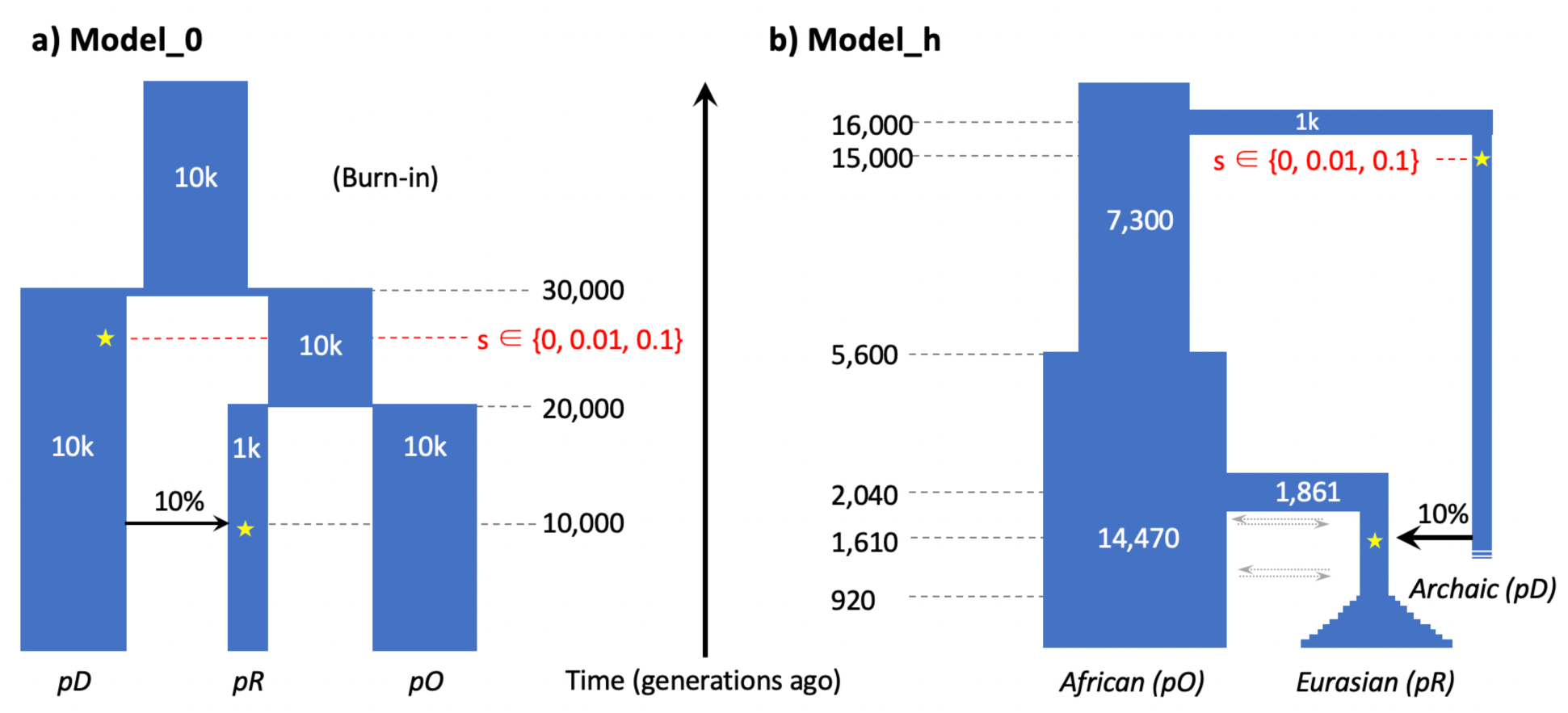
Simulated demographic models. Going forwards in time, after a burn-in period of 10*N generations (100k generations for Model_0 and 73k for Model_h), the ancestral population diverged into two subpopulations, pD and the ancestral population of pO and pR. The second population split results in pR and pO. Some time after the split of pO and pR, a single pulse of admixture occurred such that 10% of the ancestry of the recipient population (pR) came from the donor population (pD). In the presence of positive selection, a mutation was introduced at a single site in an exon for all genomes in the donor population (selection coefficient at 0.01 or 0.1). In the neutral and deleterious simulations, the selection coefficient of this particular mutation is set to 0. Except for the Neutral simulations, all other simulations contain deleterious mutations drawn from a gamma distribution of selective effects with shape parameter of 0.186 and average selection coefficient at −0.01315.

Kim *et al.*^15^reported that a long-term population contraction can greatly influence the dynamics of introgression, and that a prolonged bottleneck in the recipient population leads to a drastic increase of introgressed ancestry when the deleterious mutations are recessive. Thus, we use Model_0 as a general model to examine the robustness of the summary statistics when the heterosis effect from recessive deleterious variants is maximized. In contrast, Model_h serves as a comparison to evaluate the behavior of the summary statistics under a realistic demography for human populations.

We introduced mutations in the simulations that could have one of four different effects on fitness: 1) “Neutral”: all mutations being neutral (*s*=0); 2) “Deleterious”: recessive deleterious mutations present in the populations, drawn from a gamma distribution of fitness effect (DFE) with a shape parameter of 0.186 and average selection coefficient of −0.01315(see Kim *et al*.^47^), as well as a 2.31:1 ratio^48^ of nonsynonymous to synonymous mutations; 3) “Mild-Pos”: the Deleterious model with an adaptive mutation with milder strength of positive selection (*s*=0.01) introduced in pD (donor population) after the initial pD-pO split; 4) “Strong-Pos”: the Deleterious model with an adaptive mutation with stronger strength of positive selection (*s*=0.1) introduced in pD after the initial split.

All simulated sequences have a length of 5MB, with a genic structure that includes exons, introns, and intergenic regions. Under each model described above, we simulated 1) a 5MB region with the genic structure of a window in the human genome^49^ that has the highest density of exons (chr11:62.3-67.3MB; referred to as “*Chr11max*”; Supp. Fig. 1; also see Methods); 2) 5MB regions surrounding the previously identified adaptive introgression candidate regions in modern humans (Supp. Table 1), with the candidate region centered at approximately 2.5MB. To observe the effect of recombination rate (*r*) on the heterosis effect, we simulated recombination rate using either: 1) the realistic recombination rate map for humans^50^ inferred from linkage disequilibrium (LD) patterns^51^ and the known rates from pedigree studies^52,53^; or 2) an uniform low recombination rate at 1e-9 per base pair per generation.

For each simulation replicate, we computed the summary statistics for detecting adaptive introgression for non-overlapping 50kb windows throughout the simulated segment using a customized Python script. A full list of the AI summary statistics used in our study can be found in Table 1. We also recorded the ancestry in the recipient population that originated from the donor population using the tree sequence file generated from SLiM, and reconstructed the information using pyslim^54^ and msprime^55^ modules in Python3, which was referred to as “introgressed ancestry” or pI^43^. Throughout the text, we refer to pD as the donor population, representing an archaic hominin group; and pO as the outgroup, representing an African non-admixed population; and pR as recipient population, representing a non-African admixed population.

**Table 1:** Summary statistics informative about AI examined in this study.

| Statistic | Definition | Reference |
| --- | --- | --- |
| pI | Ancestry in the recipient population introgressed from the donor population. This measurement is directly tracked in simulations using tree sequences. | Kim *et al.* 2018; Kelleher *et al.* 2016 |
| RD | Average ratio of sequence divergence between an individual from the recipient and an individual from the donor population, and the divergence between an individual from the outgroup and an individual from the donor population | Racimo *et al.* 2017 |
| D | Patterson's D statistic, which measures the excess allele sharing between the recipient and donor population than between the recipient and an outgroup population that is unadmixed. | Green *et al.* 2010 |
| fD | A statistic that measures the excess allele sharing while controlling for local variation in the recipient population | Martin *et al.* 2015 |
| U20/U50/U80 | Number of uniquely shared alleles between the recipient and donor population that are of frequency < 1% in the outgroup, 100% in the donor, and more than 20/50/80% in the recipient population | Racimo *et al.* 2017 |
| Q90/Q95 | 90/95% quantile of the distribution of derived allele frequencies in the recipient population, that are of frequency below 1% in the outgroup and 100% in the donor population. | Racimo *et al.* 2017 |
| Heterozygosity | Expected heterozygosity in the recipient population measured by the mean of $2 \cdot p \cdot (1-p)$, with p being the frequency of any given allele in the recipient population | Crow *et al.* 1970 |

### Recessive deleterious variants affect the summary statistics used to detect AI

We first sought to understand how the presence of recessive deleterious variants affects the distribution of the AI summary statistics listed in Table 1. To maximize the heterosis effect, here we simulated the genic structure of the “*Chr11Max*” genomic region with a uniformly low recombination rate (r=1e-9) under the Model_0 demography.

Fig. 3 shows the distribution of one of the summary statistics, U80 in non-overlapping 50kb windows. U80 captures the number of high-frequency introgressed-derived alleles in the recipient population. Under the scenario where all mutations are neutral, we expect the dynamics of introgressed-derived alleles to be influenced simply by gene flow and other subsequent neutral processes. With a small pulse of admixture, only a small fraction of the introgressed alleles is expected to drift to high frequencies, which is reflected by the low to zero U80 allele count in the distribution of U80 under the Neutral simulations (Fig. 3a). However, in the presence of recessive deleterious variants, the count of U80 alleles becomes elevated in all genomic windows. This pattern is illustrated by the substantially increased mean and variance in the distribution, in contrast to the Neutral comparison (Fig. 3a). In cases of adaptive introgression where a beneficial mutation is introduced in the donor population prior to admixture (Fig. 3c-d), a notable increase of the mean and variance of U80 is also observed. Therefore, the signatures of adaptive introgression and the heterosis effect due to deleterious mutations are similar, but AI leads to a more pronounced peak at the beneficial mutation. Additionally, an adaptive mutation elevates the range of summary statistics in the flanking region, and the length of the region under its influence positively correlates with the strength of selection. However, when the elevation in U80 is due to recessive deleterious mutations, there is a slight but consistent upward shift across the entire region.

**Figure 3:**
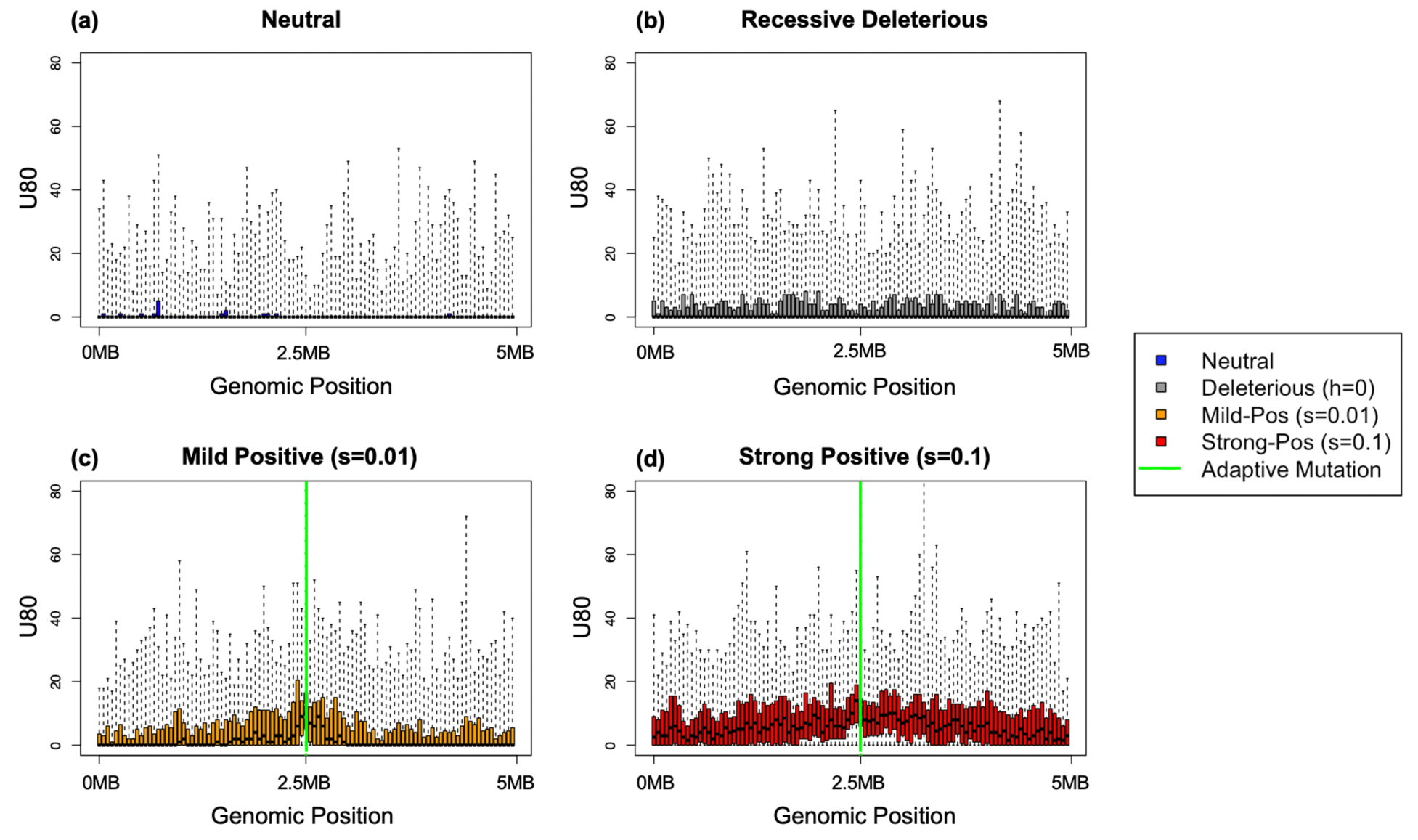
U80 statistics range under Model_0. Panels a-d respectively show the distributions of U80 statistics in 50kb windows across the 5MB region in Chr11 when mutations are neutral *(panel a)*, recessive and deleterious *(panel b)*, mildly beneficial *(panel c)*, and highly beneficial *(panel d)*. Recombination was simulated at a uniform rate of 1e-9. The adaptive mutations in the latter two models are introduced in a window in the middle of the region (2.5MB), indicated by the green solid line. Panel b, c and d also carry deleterious mutations drawn from a gamma DFE distribution. The plot shows the interquartile distributions of U80 in boxes, with whiskers extending to all data points.

We next examined the distribution of other summary statistics under the four fitness scenarios (Supp. Fig. 2), and observed similar patterns as for U80. These findings indicate that consistent with what Kim *et al.* observed for introgressed ancestry, deleterious variations can generate similar patterns as adaptive introgression in the absence of beneficial alleles and local adaptation.

To better understand the spatial patterns of variation across the simulated region, we visualized the haplotypes^56^ in a 100kb window in the middle of the segment containing the adaptive mutation when applicable (Fig. 4). The haplotype left by recessive deleterious mutations (Fig. 4a) and a legitimate adaptive mutation (Fig. 4b) differ in structure. Interestingly, both scenarios lead to higher haplotype homozygosity in the recipient population. However, in the AI scenario (Fig. 4b), the haplotypes from the donor and recipient populations are more alike to each other (i.e. the number of differences between the donor haplotype and the introgressed haplotype is smaller, shown in the right panels of Fig. 4) than under the Recessive Deleterious scenario.

**Figure 4:**
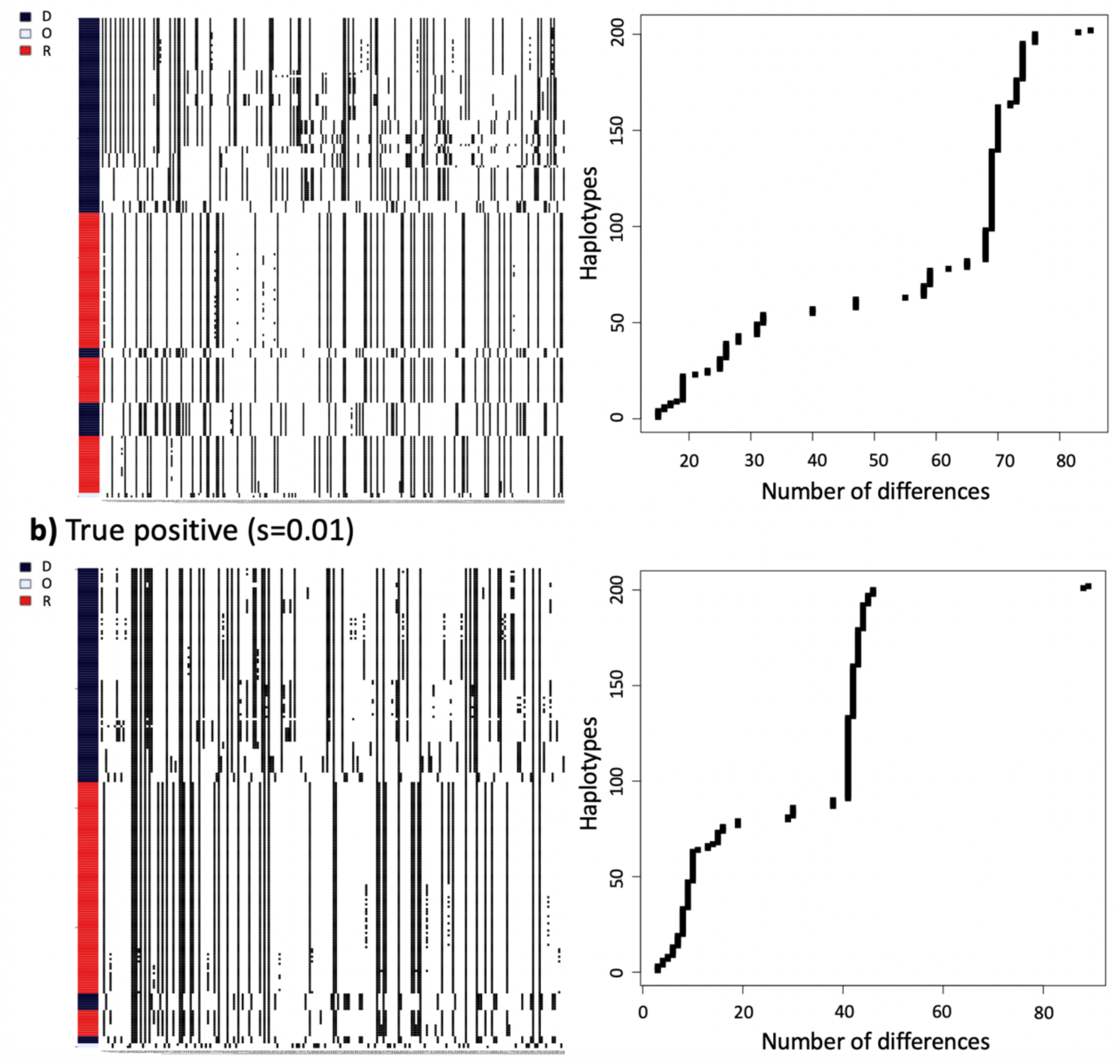
Haplotype patterns at 100kb window surrounding the adaptive allele. For each type of simulation, we sampled 100 haplotypes (rows in the heatmap) in the middle 100kb region of the Chr11Max segment each from the donor and recipient populations, and 2 haplotypes from the outgroup population (Model_0 simulations, with uniform recombination rate at 1e-9). We order the haplotypes the clusters by decreasing similarities to the donor population haplotypes (See Methods). The panels next to the heatmaps label the donors (pD, in black), recipients (pR, in red) and individuals from the outgroup population (pO, in blue). The right-hand side of panels a and b are the number of differences between the donor haplotypes and the individuals in the recipient population sorted by decreasing similarity.

### Deleterious mutations increase the false positive rate for AI detection

To quantify the extent to which deleterious mutations can give false evidence of adaptive introgression, we used the neutral distribution of summary statistics in each 50kb window across the large 5Mb segment to define the critical values for a test of adaptive introgression. We define the critical value as the most extreme 5% quantile value grouping all windows from neutral simulations together.

For the recessive deleterious model, we obtain the proportion of simulations (200 replicates) per window that exceeds the critical value under the neutral model, and define this proportion as the false positive rate (FPR), as no true adaptive mutations are present. Similarly, we define the true positive rate (TPR) for the mild- and strong-positive selection models as the per-window proportion of simulations exceeding the critical value, where the critical value is again defined from the neutral model. Fig. 5 shows the neutral critical value and the transformed true/false positive rates in U80 and Divergence Ratio (RD) statistics under the simulation setting described in the above section. The TPR/FPR distribution for other summary statistics can be found in Supp. Fig. 3. The neutral model simulations have FPRs around 5%, by definition. In contrast, the recessive deleterious simulations show elevated FPRs in most windows for both statistics (8.62-34.48% for RD; 3.45-22.41% for U80). The high FPRs are not negligible, as the identification of AI in empirical data relies on looking for outliers in summary statistics when the presence and location of the adaptive mutation is unknown. Deleterious variation is also more common in human genomes than adaptive variation^34^, which may further compound this effect.

**Figure 5:**
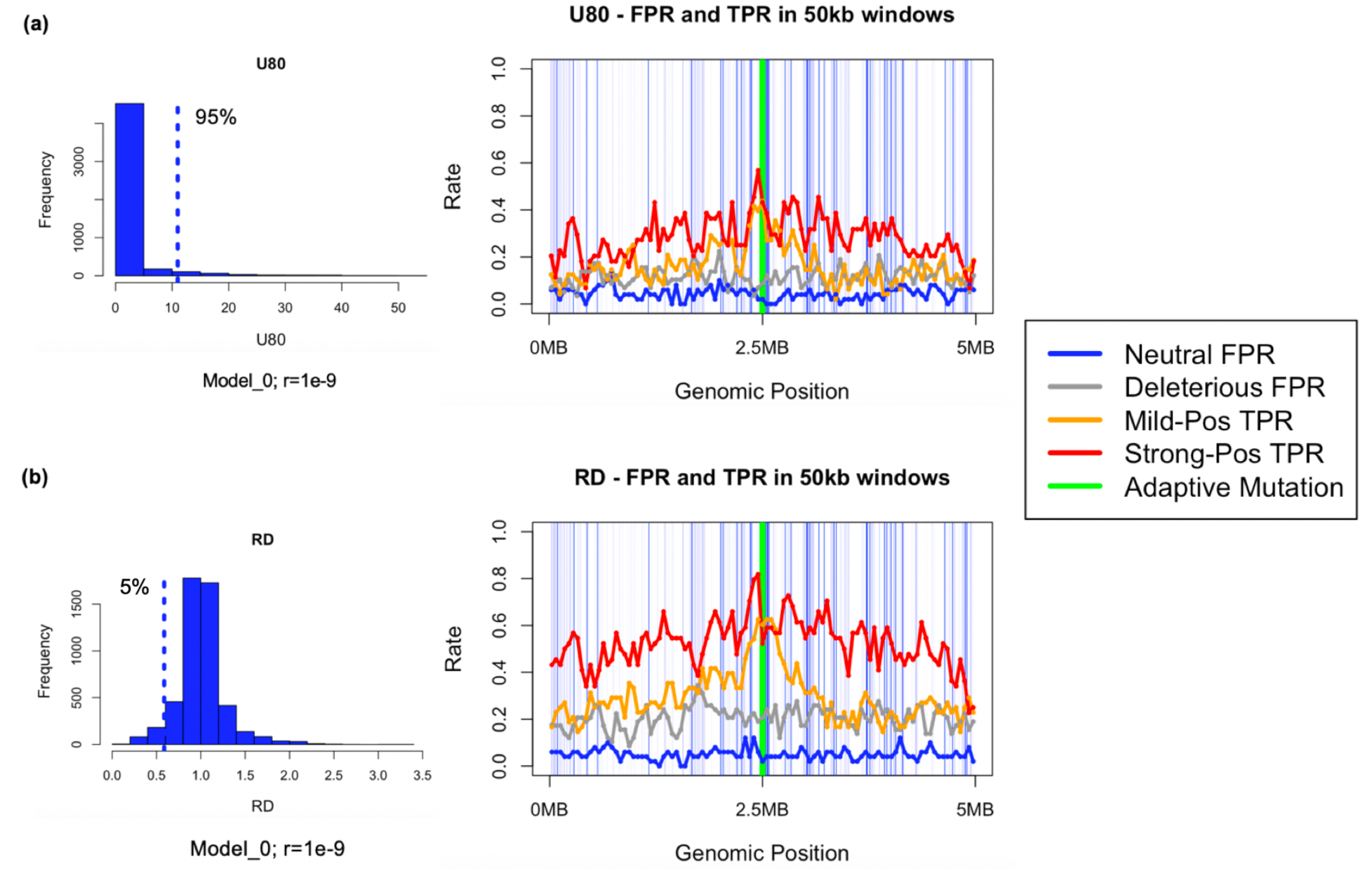
Distributions and True/False Positive Rates of U80 and RD Statistics. This figure shows the critical value (dotted line) used to compute the False/True Positive Rates for two summaries of the data: U80 and RD (left hand side of panels a and b). The right-hand side of panels a and b show the False Positive Rates (under the neutral and deleterious models) and the True Positive Rates (under the models with positive selection) for each 50kb window in a region of 5MB. For the simulations, red, orange, blue and black represent Strong-Pos, Mild-Pos, Neutral, and Deleterious respectively. The light blue lines in the mid-panels illustrate the exons where new mutations can arise, and the green solid line represents the window where the adaptive mutation occurred. The simulations ran under Model_0 using the genic structure of the Chr11Max region, using a uniform low recombination rate of 1e-9.

To further understand how demographic history and recombination influence the FPR/TPR of the tests for AI, we simulated the “*Chr11Max*” 5MB segment (see Simulations section) using the human demographic model (Model_h), and realistic estimates of recombination rate in this region (referred to as r=hg19 in Table 2). We summarized the FPRs and TPRs of a subset of statistics (pI, RD, U80, Q95) under these scenarios in Table 2 (also see Supp. Fig. 4-6). We observed that simulations with low recombination rate with higher mean FPRs using these statistics. Moreover, the standard deviation of the statistics – which is an informative signature of adaptive introgression – increases when the realistic recombination rates are applied (average recombination rate higher than 1e-9).

**Table 2:** Summary of the TPR/FPR under different models.

| Simulation Scenario | Statistics | Mean of FPR in Deleterious Model | SD of FPR in Deleterious Model | Focal Window TPR in Mild-Pos Model | Focal Window TPR in Strong-Pos Model |
| --- | --- | --- | --- | --- | --- |
| Model_0;<br>*Chr11Max*;<br>$r=1e-9$ | pI | 0.354 | 0.047 | 0.900 | 1.000 |
|  | RD | 0.204 | 0.048 | 0.521 | 0.569 |
|  | U80 | 0.117 | 0.038 | 0.375 | 0.431 |
|  | Q95 | 0.437 | 0.051 | 0.875 | 1.000 |
| Model_0,<br>*Chr11Max*;<br>$r=hg19$ | pI | 0.229 | 0.086 | 0.885 | 1.000 |
|  | RD | 0.134 | 0.061 | 0.577 | 0.648 |
|  | U80 | 0.090 | 0.045 | 0.442 | 0.5 |
|  | Q95 | 0.121 | 0.034 | 0.637 | 0.752 |
| Model_h;<br>*Chr11Max*;<br>$r= hg19$ | pI | 0.087 | 0.108 | 0.967 | 1.000 |
|  | RD | 0.098 | 0.117 | 1.000 | 0.654 |
|  | U80 | 0.022 | 0.049 | 0.667 | 0.500 |
|  | Q95 | 0.099 | 0.120 | 1.000 | 0.933 |

For the deleterious model, we computed the false positive rates (FPRs) in 50kb non-overlapping windows using the most-extreme 5% value from the neutral distribution as critical value, and show the mean FPR in the third column. For the adaptive introgression models (Mild-Pos and Strong Pos), we computed the true positive rates (TPRs) using the same neutral cutoff value in all windows, and show the true positive rate in the window that contains the adaptive mutation ("Focal TPR"). Note that a properly calibrated null model should have a FPR of 0.05.

Altogether, recessive deleterious variants contribute to a higher false positive rate for AI detection in all summary statistics examined. Some statistics appear to be more vulnerable than others, with pI, RD, U stats and Q stats being most affected. Low recombination rates amplify the heterosis effect that mimics the AI signature, while the modern human demography (Model_h) results in fewer false positives than Model_0 in general which has a relatively long-term contraction in the recipient population.

### Summary statistics are powerful to detect AI but not at localizing the adaptive allele

We next evaluated the power of these summary statistics at detecting true AI. The TPRs across the genomic region is not uniform (Supp. Fig. 3-5). On average, the TPRs are close to or higher than the FPRs in corresponding windows. In the focal windows containing the adaptive mutation, the TPRs are especially distinguishable from the neutral and deleterious models because the adaptive models show a distinct peak. This shows that the summary statistics have high statistical power at detecting a true AI signal, as they reject the null hypothesis more often in true positives (up to 100%; Table 2).

The mean TPR is parameter-dependent like the FPR, in that it increases with selection strength and it decreases with recombination rate. However, it should be noted that under very strong positive selection (Strong-Pos model), the TPRs are high across longer flanking regions, resulting in the focal window not standing out from the background because the region affected by positive selection is larger. In the weaker positive selection (Mild-Pos) model, the focal window stands out with respect to the background windows because the background windows reject the null less often. Therefore, localizing the adaptive allele within the entire segment becomes less accurate with increasing strength of selection on the truly adaptive allele (Fig. 5, Supp. Fig. 3-5).

### Deleterious mutations have a limited effect on candidates for adaptive introgression in humans

Next, we sought to systematically assess whether the changes in AI summary statistics caused by recessive deleterious variants could lead to false detection of AI candidate regions in humans. This is an important consideration because these regions were detected as unusual either in comparison to the rest of the genome or under demographic models that assumed all mutations were neutral. Thus, the null models did not include deleterious variation and it remains unclear whether deleterious variation could provide an alternate mechanism for the observed patterns.

We extracted 5MB sequences surrounding 26 previously identified AI regions^9,11,13–18,20,22,57–59^ (Supp. Table 1) using the distribution of the recombination rates^52^ and genic structures^49^ in these regions. For each candidate region, we ran 200 simulation replicates under a more realistic human demography (Model_h), using the recombination rates and exon distribution from these regions. We simulated a recipient population representing a non-African population (pR), an outgroup population (pO) representing Africans, and an archaic donor population (pD). In addition, we simulated under two models (the Neutral and Deleterious models) to compute the FPRs on summary statistics within each 50kb window in each of the 5MB regions representing the AI candidate gene-regions.

To compute the false positive rate due to deleterious mutations, we use the neutral simulations to define the critical values for each test statistic. We use the simulations with recessive deleterious mutations as the test datasets to examine the false positive distribution. As described previously, the FPR represents the proportion of simulations for a given statistic in a 50kb window in a candidate gene that are as extreme or more extreme than the 5% neutral critical value.

Overall, we find that most statistics do not have extremely elevated false-positive rates across most of the gene-regions in the presence of deleterious mutations (Supp. Fig. 6). The D statistic, however, is a notable exception showing a higher FPR across all candidates. This is rather unsurprising because, although the D statistic is powerful at detecting genome-wide excess of shared derived alleles between groups (a metric indicating admixture), studies have shown its limitations and reduced reliability for inferring local ancestry using small genomic regions^26^ (50kb windows). The fD statistic, on the other hand, is powerful at detecting introgression at localized loci, and does not show unusually high FPR for all candidate regions.

Notably, with the exception of two simulated regions (representing the regions of *HLA* and *HYLA2*, Fig. 6), we find that the FPR is well-controlled in the other 24 simulated AI candidate regions (Supp. Fig. 6). Here, we show the FPRs for the *EPAS1* and the *BNC2*-like regions (Fig. 6) since these two regions have similar recombination rates, exon density and FPRs as the other AI regions considered here. Other than the D statistics discussed above, the rest of the summary statistics show an average of FPR around or less than 5%. In particular, the Q and U statistics appear to be the most robust against false positives from deleterious mutations. In contrast, *HLA-A, HLA-B*, and *HLA-C* genes (referred to as “*HLA*” in this work), and a segment on chromosome 3 that contains *HYAL2* gene show elevated FPRs on nearly all statistics.

**Figure 6:**
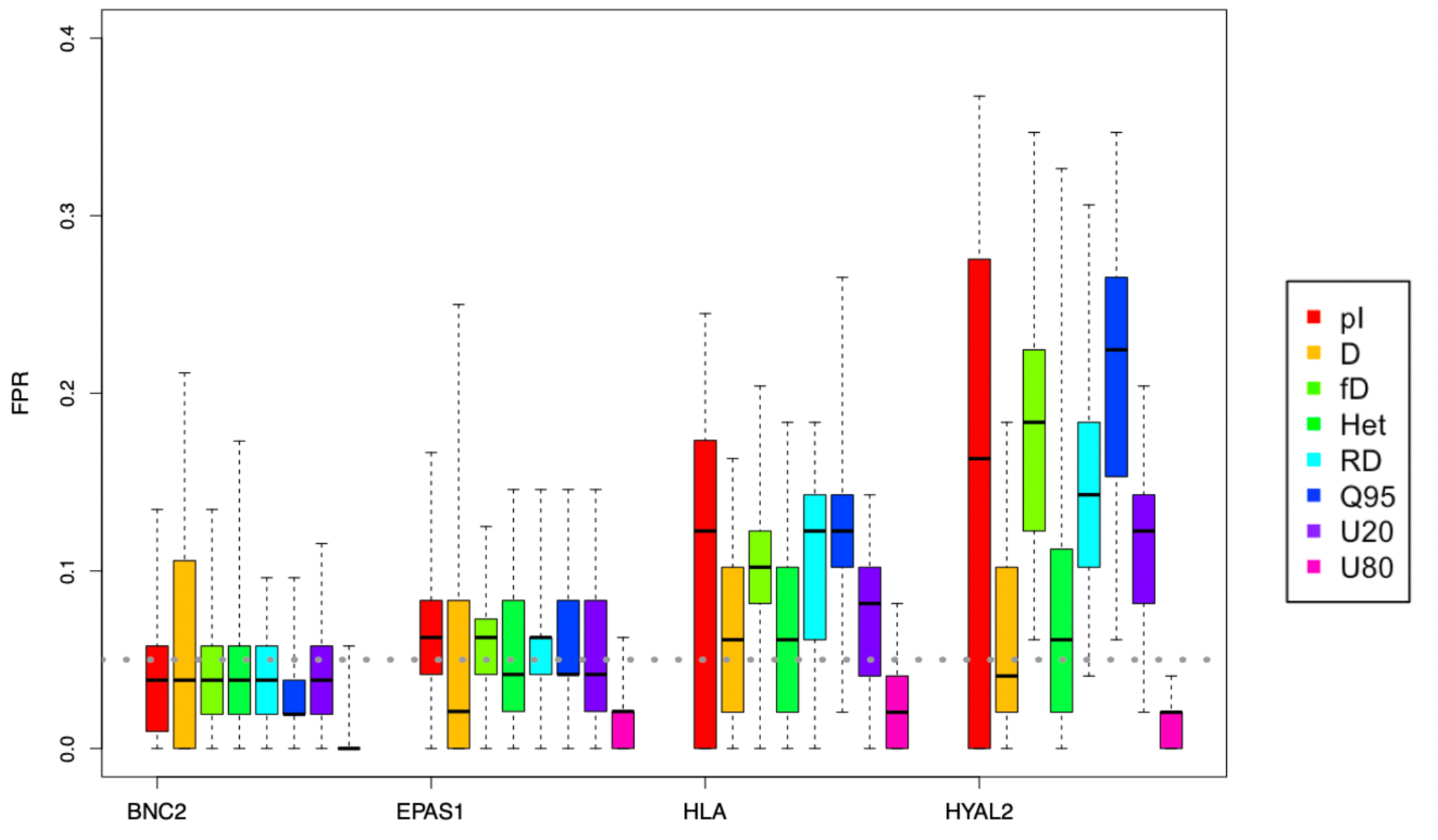
False Positive Rates for summary statistics from human AI candidate regions. The summary statistics are obtained from simulations under Neutral and Deleterious mutation models using human demography, Model_h. The recombination rates and exon density reflect the four regions in the human genome harboring BNC2, EPAS1, HLA and HYAL2. The FPR (y-axis) is computed assuming a neutral null model and represents the proportion of simulations replicates under the Deleterious model that are called significant for adaptive introgression. The HLA and HYLA2-like regions result in the highest FPRs, while the EPAS1 and BNC2-like regions have similar FPRs as the other regions simulated.

### High exon density and low recombination rate leads to deleterious mutations mimicking AI in humans

To understand why the *HYAL2* and *HLA* genes exhibit higher false positive rates in the presence of recessive deleterious variants, we considered possible sources of false positives. Firstly, we wanted to know whether population growth in humans was a contributing factor. Secondly, since deleterious recessive mutations are much more likely to occur only in exons, we looked at the distribution of exon density in windows of 5MB across the genome to ask whether *HYAL2* and *HLA* are outliers with respect to this distribution. In addition, we computed the recombination rate for each 5MB window across the genome to determine whether these two genes were outliers with respect to recombination rate.

We first simulated the four genes in Fig. 6 under four different scenarios of population size changes (Supp. Fig. 7). We find that the extent of population growth does not play a determining role on the FPRs in AI detection since in our simulations. Specifically, the outlier regions such as *HYAL2* and *HLA*, continue to have higher FPRs across the different growth scenarios. Growth (eg. “*Growth 2*” and “*Growth 4*” in Supp. Fig. 7 where the population size at the end generation is more than 70-fold larger than the initial size) slightly intensifies the already high FPRs in these two genes (Supp. Fig. 8), which can be explained by an increase in the efficacy of selection when the effective population size is large^60,61^. The other two simulated regions (representing the *BNC2* and *EPAS1* regions) do not exhibit increased FPRs in the presence of population growth.

We next explored how changes in recombination rate impact the FPRs for the summary statistics. By applying a uniformly low or high recombination rate to the simulations under Model_h (Supp. Fig. 9), we observed that a high recombination rate can substantially reduce the FPRs to nominal levels (around 0.05) on all statistics in all genes. Conversely, a uniformly low recombination rate does not necessarily increase the FPRs in most statistics when we simulate regions like *BNC2* and *EPAS1*, except for the D statistics.

Finally, we computed the mean recombination rates at 5MB windows across the human genome (Hg19), and demonstrate that *HYAL2* and *HLA* regions are indeed outliers in terms of both exon density and low recombination rate (Fig. 7). We therefore conclude that the high susceptibility to false detection of AI in some genomic regions, is due to a combination of high exon density and low recombination rate. This also explains why the confounding effect of heterosis is limited for the majority of the human AI candidate gene regions simulated here (Supp. Fig. 10).

**Figure 7:**
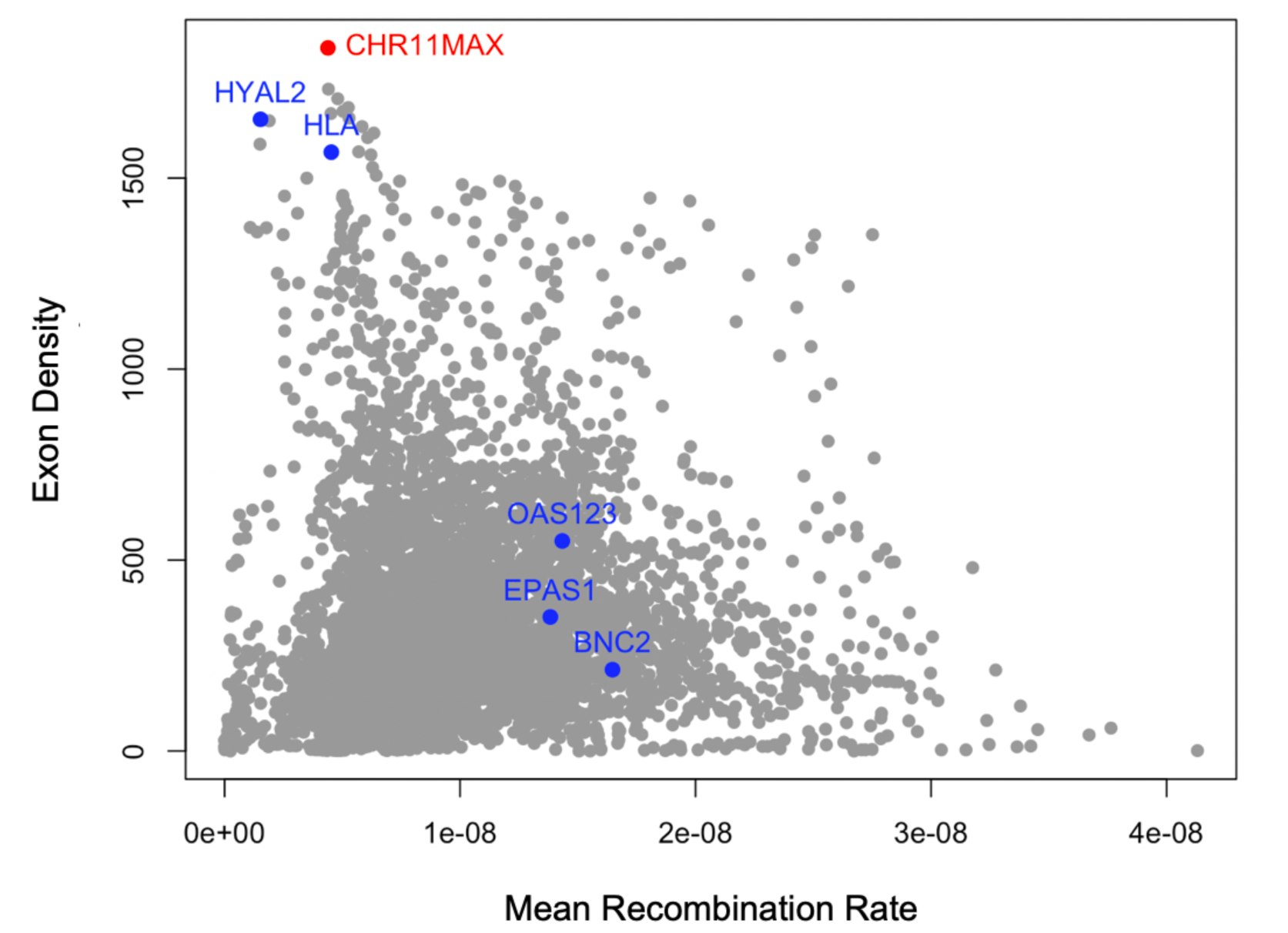
Human whole-genome exon density and mean recombination rate. This figure shows the relationship between the exon density and mean recombination rate in non-overlapping 5MB windows across the human genome (hg19). The blue points highlight the regions of AI candidate genes mentioned in the main text, including the outliers (HYAL2, HLA), and the typical ones (EPAS1, OAS cluster, BNC2). The red point represents the “Chr11max” region mentioned in earlier sections.

### Null model with deleterious variation reduces the number of statistically significant AI candidates

Lastly, we asked whether a null model that accounts for the recessive deleterious variants can be more informative and reliable in AI detection than a traditional null model that assumes selective neutrality. To do this, we calculated the empirical values of the summary statistics from the AI candidate genes from the 1000 Genomes Project dataset^62^using one of the archaic humans (Altai Neanderthal^5^ or Denisovan^8^) as the donor population, and the Yorubans (YRI) as the outgroup population. We computed their p-values using the statistical distributions from the simulations under two different (Neutral or Deleterious) null models. Given that our deleterious null model assumes all deleterious mutations are recessive (h=0), it maximizes the impact of false positives due to deleterious mutations. Thus, if the candidate genes still stand out as being statistically significant in this extreme null model, the AI signal is robust to confounding from the heterosis effect.

We calculated the critical values for all summary statistics using the most extreme 5% tail values under the two null models, and computed the p-values of the empirical data points for the statistics. Among the four genes we use as examples (Supp. Fig. 11), the “outlier” genes (*HLA* region and *HYAL2*) on average have higher p-values under the deleterious null models than under the neutral null model. This trend is reflected by the points falling mostly above the diagonal in Supp. Fig. 11. Since higher p-value implies that we cannot reject the null model, this change in the p-values indicates that the deleterious null models are more conservative at AI inference. Note, that for the two “typical” AI genes, the p-values fall along the diagonal (Supp. Fig. 11), suggesting that a null model with and without deleterious mutations yield similar results.

We also examined the number of 50kb-windows that fell in the extreme 5% tail of the neutral distributions, as well as that number from the deleterious distributions. The difference between the two numbers equals the number of window hits that are significant in the neutral null models but failed to reach significance in the deleterious null models (Fig. 8, Supp. Fig. 12). The positive values, highlighted in the gray-shaded area in the corresponding figures and colored by population, imply the deleterious null model is more conservative for a given statistic. If an AI candidate region shows points above zero for most of the summary statistics, such candidate region is likely prone to false positives due to the heterosis effect, and the validity of adaptive introgression on this region requires further investigation.

**Figure 8.**
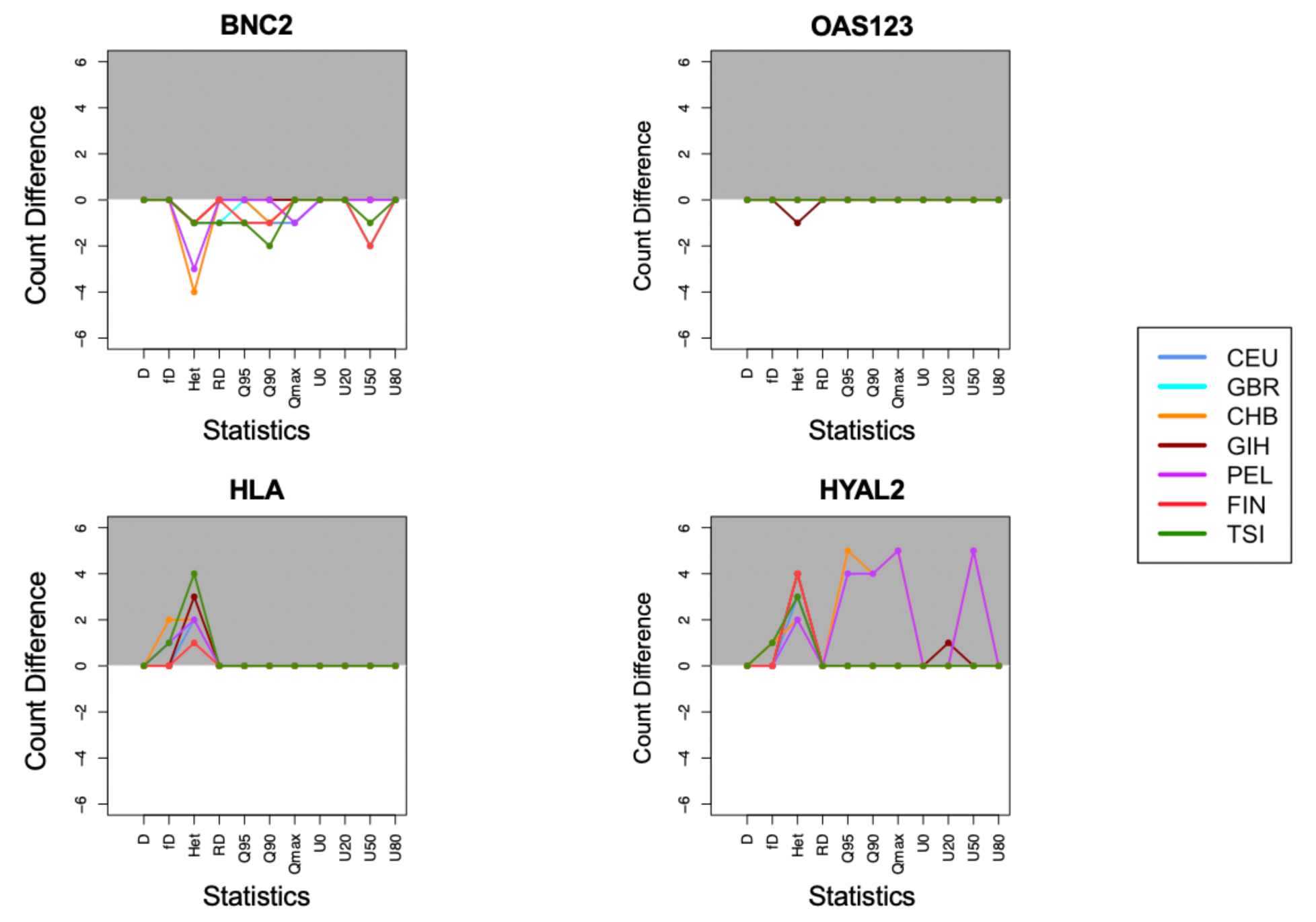
Significant hits number change in candidate genes between different null models. We compared the difference in the number of significant hits (windows with p-value <0.05) predicted by neutral and deleterious null models. Each point represents that difference in number (y-axis, Neutral significance number – Deleterious significance number) in its corresponding statistics (x-axis). The genes with multiple points above y-axis value 0 are highlighted in the gray-shaded area, indicating the deleterious null models predict fewer window hits being significant for given statistics, which implies potential false positives from neutral null models. The OAS gene cluster (“OAS123”) - is shown here instead of EPAS1 because the AI signal in EPAS1 is not shown in any of the 1000 Genomes populations.

Promisingly, we find that most of the candidate regions (24/26) show similar p-values on most, if not all, of the statistics regardless of whether a null model with deleterious mutations or neutral mutations is used. This observation further confirms the conclusion from the previous section, that the distribution of recessive deleterious variants has a limited impact on the detection of adaptive introgression in modern humans.

As shown in our analysis, a combination of high exon density and low recombination rate contribute to the high FPRs in the *HLA* and *HYAL2* genes, with both showing a reduced signature of adaptive introgression under a deleterious null model. This suggests that these regions potentially may not be adaptively introgressed, in contrast to previous findings^9,11,14,22^. The *HLA* cluster on Chromosome 6 (*HLA-A, HLA-B, HLA-C*) is one of the most crucial immune-response genes, and is known for its high level of genetic variation and variability across populations^15,63,64^. Because of the complexity of population genetics processes (e.g. balancing selection) that act on *HLA*, further work is required to understand whether deleterious mutations or other types of selection can lead to the behavior of summaries of genetic variation at this region. For example, the high FPRs assume a null deleterious model which does not explain the high levels of heterozygosity (Supp. Fig. 13) at this locus, so the evolutionary processes acting for this region are more complex than the null model assumed. However, in general integrating recessive deleterious mutations into the modeling framework will improve the robustness of adaptive introgression signals.

## Discussion

This work represents one of the first comprehensive efforts to consider the influence of negative selection in the detection of adaptive introgression. Specifically, we systematically examined whether deleterious recessive variants carried by populations prior to admixture can affect the robustness of signals in summary statistics that have been shown to be informative about adaptive introgression.

Our work demonstrates through extensive simulations that a heterosis effect caused by recessive deleterious variants private to source populations can resemble the signal of adaptive introgression, which leads to a higher number of false positives. We found that the presence of recessive deleterious mutations alone is sufficient to significantly increase the mean and variance of AI summary statistics in at least some genomic regions. These shifts in the distribution of statistics (Fig. 2) lead to a higher false positive rate for detection of adaptive introgression when we use the neutral model to define AI statistic critical values. Moreover, by examining population genomics data, we show that such effect from recessive deleterious variants are relevant for detecting AI in modern humans, and may explain a potentially spurious signal of AI in at least two AI candidate genes (*HLA* and *HYAL2*). However, the statistical signals in other candidate genes in modern humans remain strong even when accounting for recessive deleterious mutations, indicating that these candidates are unlikely to be false positives.

By testing individual evolutionary parameters in genes that show a higher magnitude of false positives than others, we attributed the stronger heterosis effect to two factors that need to present at the same time: high exon density, and low recombination rate (like in *HLA* and *HYAL2*). High exon density implies more deleterious mutations occur in a given genomic region. In most cases, the deleterious fitness effect from linked recessive variants can be disassociated from crossing over with other haplotypes within the same population. However, for certain regions where the recombination rate is unusually low, the deleterious variants will remain linked on a given haplotype. Admixture with a distantly related population will bring in haplotypes carrying non-deleterious alleles at these positions. Therefore, the introgressed ancestry at these regions will increase in the recipient population despite carrying a different set of deleterious variants, leading to the elevation of FPRs in the adaptive introgression summary statistics. This process acts in a similar manner as balancing selection, except that no beneficial mutations are involved.

We also show that the demographic history of human populations, including a change in the recipient population size, does not play a major role in affecting the false-positive rate of tests for AI. However, the nearly-exponential population growth in the recent history of modern humans may have increased the FPR in genes that are already susceptible to false-positive results due to deleterious mutations. This is likely due to the fact that a large effective population size restricts the extent of genetic drift, leading to a more prominent effect of natural selection, including the complementation of deleterious alleles via the heterosis effect. Depending on the dynamics among different types of selection, a recovery of population size after a bottleneck in the recipient population can exaggerate the heterosis effect, as demonstrated in Kim et al^43^.

Though the signals in most human AI candidate genes are unaffected by deleterious variation, the impact of deleterious variants on AI detection in general should not be omitted, and a null model that considers the influence from recessive deleterious mutations is still necessary. This is mainly because of two reasons: 1) the combination of evolutionary parameters that leads to an elevation of false-positives may occur much more commonly in other study systems. 2) Even for modern humans, the demography used in simulations is an approximation of the modern Eurasian population history, which may not represent the true evolutionary history of all non-African populations. In situations where the archaic introgression occurred in multiple pulses (e.g., Denisovan introgression in Asia^22,65^), and when the ancestral modern human populations were small, the heterosis effect from deleterious variants could have a different impact than expected from a parsimonious demography.

Here, we considered only the extreme case where deleterious variants are completely recessive (h=0). The reason for this is that we set out to determine whether deleterious variants are a concern for AI signals when this effect is maximized. Kim *et al.* already studied the effect from additive variants and observed little effect on introgressed ancestry, where the confounding effects from heterosis persisted when deleterious variants were complete or partial recessive (*hs* relationship^35^). In empirical genomic data, the distribution of dominance should be in between the two extremes. A current challenge is that the empirical values of dominance coefficients for deleterious mutations in humans remain unknown.

It is promising that the signature of AI in the vast majority of human AI candidate regions still persists even when the heterosis effect acts in its most extreme manner by assuming *h*=0. Other values of h would be unlikely to affect the conclusion that 24/26 candidates are robust to confounding by deleterious mutations. The *HLA* and *HYAL2* genes are outliers in terms of their exon density and recombination rate which accentuates the effect of heterosis, and further increases the probability of observing extreme summary statistics in a model with recessive deleterious mutations. In general, our present study shows that if deleterious mutations are completely recessive, they can account for most of the AI signatures in these two genes. However, if deleterious mutations are only partially recessive, then it is possible that, by themselves, deleterious mutations cannot account for these putative signals of adaptive introgression. In such a scenario, true AI would be required to explain the data. It is also worth mentioning that *HLA* gene exhibits complex patterns of genetic variation. We find that even a conservative deleterious recessive model cannot generate the levels of heterozygosity observed for *HLA* (Supp. Fig. 13), and more work is warranted to determined what actual evolutionary processes are acting in this region.

Our study demonstrates from multiple angles that the presence of recessive deleterious variants in populations can sometimes generate similar statistical signals as adaptive introgression in the absence of any beneficial alleles. Although more conservative, it results in inferences that are more robust compared to a neutral null model. We should bear in mind that the overall robustness of the AI signals in modern humans may be attributed to many factors including the unique genic structure, and the difference in AI signature contributed by the distribution of recessive deleterious variants is still not negligible. We therefore strongly encourage future AI studies to use a null model that incorporates a realistic distribution of fitness effect for deleterious variants, recessive or partially recessive, to minimize false positives. This approach is particularly relevant for studying organisms that have more compact genomic structures, and/or different demographic histories that may accelerate the dynamics of the heterosis effect after introgression.

## Acknowledgement

This work was supported by NIH grant R35GM119856 (to K.E.L) and E.H.S was supported by NIH grant 1R35GM128946-01 and NSF grant 7378-#1557151. The authors thank their colleagues from the Lohmueller Lab at UCLA and the Huerta-Sanchez Lab at Brown University for helpful discussions during the development of this study. We also thank Dr. Fernando Racimo at University of Copenhagen, Denmark for kindly sharing sample code for computing AI summary statistics.

## Methods

### Forward Simulations

We used the software SLiM (version 3.2.0)^44^ throughout this work for the simulations. All mutations that became fixed in all population by the end of the simulations were disregarded from downstream calculation of summary statistics. We chose to use the default calculation of the fitness effect for recessive deleterious mutations (h=0).

We considered three types of simulations, distinguished by the types of mutations they carry: 1) neutral simulations (“*Neutral*”): only neutral mutations are introduced (*s*=0); 2) deleterious simulations (“*Deleterious*”): in addition to the neutral mutations, we introduced deleterious mutations that are recessive (*h*=0), with a distribution of fitness effect drawn from a gamma distribution previously estimated (shape parameter = 0.186; mean selection coefficient = −0.01314). The deleterious mutations can only accumulate at exon regions, with a ratio of nonsynonymous to synonymous mutations at 2.31:1; 3) positive selection simulations: this type of simulation is subdivided into two types depending on the selection strength of the beneficial mutation introduced (“*Mild-pos*”, *s*=0.01; “*Strong-pos*”, *s*=0.1). This simulation type carries the same distribution of neutral and deleterious mutations as in group 2, while we also introduced a nonsynonymous beneficial mutation in an exon approximately in the middle of the 5MB segment in all haplotypes from the donor population after the initial split between the donor and outgroup populations. Therefore, after the single pulse of admixture from the donor to the recipient populations, at least one haplotype from the recipient population should carry the beneficial mutation that arose from the donor population. Simulation replicates where the beneficial mutation was lost from the recipient population before the end of the simulation were discarded. We obtained 200 replicates for each unique combination of simulation type and genomic structure.

We also scaled the simulation parameters by a scaling factor of *c* (*c*=5) to increase computational efficiency. The population size thus was rescaled to *N/c*, all generation times to *t/c*, selection coefficient to *s*\**c*, mutation rate to *μ*\**c*, and the recombination rate also at *r*\**c* (approximation from 0.5(1-(1-2^*r*^)^*c*^) for small *r* and small *c*). Other evolutionary parameters remain the same before and after rescaling.

### Simulations with modern human genomic structure

Unless specified separately, all simulations in SLiM from this study use genic structure from modern human genome build GRCh37/hg19. We fix the simulation segment length at 5MB, and used the exon ranges defined by the GENCODE v.14 annotations^49^ and the sex-averaged recombination map by Kong *et al.*^52^ averaged over a 10kb scale. The per base pair mutation rate was fixed at 1.5*10^−8^. For comparison purposes, we also applied a uniform recombination rates at 1.0*10^-8^ and 1.0*10^-9^ as specified in the main text.

For simulations mimicking specific adaptive introgression candidate genes, we identified the genomic coordinates using the original studies that identified the AI candidate genes (Supp. Table 1), and extracted their flanking regions upstream and downstream of the gene region to a total length of 5MB, with the gene region positioned in the center. We then used the recombination map and the distribution of genomic segments mentioned above in the simulations.

### Computing the exon density across the human genome

To tabulate exon density across the genome, we scanned the 22 autosomes of human genome using a sliding window of 5MB with step size of 1kb, and counted the number of exons per 5MB window. We defined “exon density” as the total number of exons/window. We extracted the coordinates of the window that has the highest exon density, and designated it as the “*Chr11max”* region (hg19 Chr11:62.3-67.3MB).

### Summary statistics for the detection of adaptive introgression

We directly tracked the introgression-derived ancestry (*pI*) in the recipient population from SLiM program by tracking the tree sequences across the simulated segments. Therefore, the introgressed ancestry calculated from this study is the true proportion of ancestry. The amount of *pI* was recovered from the tree sequence file generated from SLiM using a custom python script using *pyslim* module^54^.

For the other summary statistics that capture the signature of adaptive introgression (Table 1), we used a custom Python script to extract the sampled haplotype matrices that are in ms format from the SLiM output (100 haplotype samples per population), and filled in the non-segregating ancestral alleles to match the size of the haplotype matrices from the donor, recipient, and outgroup populations respectively. We calculated the summary statistics at non-overlapping 50kb windows using the same python script pipeline for each simulation replicate.

### Summary statistics for non-African modern human populations

We calculated a variety of AI summary statistics using modern human genome variation data from the 1000 Genomes Project (Phase 3)^62^. To illustrate the signals of AI captured by the summary statistics from previous studies, we used all individuals from seven representative populations from Eurasia and the Americas as recipient populations (for archaic introgression). Specifically, we used Western Europeans (CEU), British (GBR), Finnish (FIN), Italians (TSI), Han Chinese (CHB), Indians (GIH), and Peruvians (PEL). We also used Yorubans (YRI) as unadmixed outgroup. For the donor population, we used the unphased, high-quality whole genome sequences from the Altai Neanderthal^5^ and/or the Altai Denisovan^8^, depending on which archaic group was identified as the AI source (Column 4 in Supp. Table 1). We referred to the coordinates of AI candidate genes listed in Supp Table 1 to identify each 5MB region centered on the candidate gene, and extracted the corresponding genomic sequences from the modern populations and their respective donor populations. We additionally removed sites in the archaic genomes that have potential quality issues (quality score < 40 and/or mapping quality < 30). If a previously identified AI gene was found to be associated with more than one archaic groups, we used only the Altai Neanderthal sequence for these cases. As we did on the simulations, the summary statistics were calculated at non-overlapping 50kb windows in the empirical data.

### Haplotype structure comparison using Haplostrips

We used the software *Haplostrips*^56^ to plot the haplotypes from the *Chr11Max* region from the simulations. The haplotype input matrix for the software was generated from SLiM by the end of one replicate of simulation, and was further truncated to include only the center 100kb region surrounding the exon where the beneficial mutation arises when applicable. We sampled 100 chromosomes from the donor and recipient populations respectively, and 2 chromosomes from the outgroup population. The software output displayed each variant within the region as a column, and each row represents a haplotype (phased from the simulation). Each population was assigned a unique color corresponding to the haplotypes from the respective population. The haplotypes were hierarchical clustered by a decrease in similarity to the sampled haplotypes from the donor population. The panels on the right-hand side representing the distribution of haplotypes in terms of the genetic distance to the donor haplotypes.

