## Supplementary Material for "Recessive deleterious variation has a limited impact on signals of adaptive introgression in human populations"

### Supplementary materials

**Supplementary Table1: Regions scanned in this study (5MB windows)**

| Coordinate (hg19) | Identifier in this study | Adaptive Introgression Gene(s) | Donor Population | Recipient population | Biological Pathway(s) | Original Report |
|---|---|---|---|---|---|---|
| 9:14-19MB | BNC2 | *BNC2* | N | Europeans | Skin pigmentation | Vernot, Akey 2014 |
| 19:32-37MB | CHR19region | *RHPN2 GPATCH1 WDR88 LRP3 SLC7A10* | N | Europeans | *multiple genes and pathways | Browning et al. 2018 |
| 3:43-48MB | CHR3region | *CCR9 LZTFL1 FYCO1 CXCR6 XCR1* | N | South Asians | *multiple genes and pathways | Browning et al. 2018 |
| 2:43-48MB | EPAS1 | *EPAS1* | D | East Asians (Tibetans) | High altitude adaptation | Huerta-Sanchez et al. 2014 |
| 11:9-14MB | GALNT18 | *GALNT18* | N | South Asians | Glycan biosynthesis | Browning et al. 2018 |
| 6:29-34MB | HLA | *HLA-A, HLA-B, HLA-C* | N, D | Eurasians, Oceanians | Immune system regulation | Abi-Rached et al. 2011 |
| 3:48-53MB | HYAL2 | *HYAL2* | N | East Asians | Cell proliferation, metabolism | Ding et al. 2014 |
| 12:50-55MB | KRT71 | *KRT71, KRT6A, KRT5* | N | Eurasians | Keratinization, ectoderm differentiation | Browning et al. 2018, Vernot, Akey 2014 |
| 16:54.5-59.5MB | NLRC5 | *NLRC5* | N | East Asians | Innate immunity sensor | Descamps et al. 2016 |
| 12:111-116MB | OAS123 | *OAS1, OAS2, OAS3* | N, D | Eurasians | Toll-like receptor signaling | Mendez et al. 2013, Sankararaman et al. 2014 |
| 15:25.5-30.5MB | OCA2 | *OCA2* | N | Eurasians | Skin pigmentation | Gittelman et al. 2016 |
| 10:93-98MB | PDE6C | *PDE6C* | N | Eurasians | Nucleotide metabolism and phototransduction | Vernot, Akey 2014 |

| | | | | | | |
|---|---|---|---|---|---|---|
| 11:117.5-122.5MB | POU2F3 | *POU2F3* | N | East Asians | Keratinocyte proliferation and differentiation | Vernot, Akey 2014 |
| 12:119-124MB | RNF34 | *RNF34* | N | Eurasians | Innate immune system | Vernot, Akey 2014 |
| 15:46-51MB | SEMA6D | *SEMA6D* | N | Eurasians | Semaphorin interaction | Vernot, Akey 2014 |
| 4:52-57MB | SGCB | *SGCB, SPATA18* | N | Eurasians | Cardiomyopathy, mitochondrial metabolism | Vernot, Akey 2014 |
| 8:85-90MB | SGCZ | *SGCZ* | N | South Asians | Cellular regulation | Browning et al. 2018 |
| 1:230-235MB | SIPA1L2 | *SIPA1L2* | N | East Asians | Neuronal signaling | Vernot, Akey 2014 |
| 17:5.06-10.06MB | SLC16A11 | *SLC16A11* | N | Native Americans | Lipid metabolism, Type 2 Diabetes | Williams et al. 2014 |
| 2:226-231MB | SLC19A3 | *SLC19A3* | N | South Asians | Vitamin digestion and absorption | Browning et al. 2018 |
| 17:16.5-21.5MB | SLC5A10 | *SLC5A10* | D | Europeans | Hexose transport and metabolism | Racimo et al. 2017 |
| 12:54.5-59.5MB | STAT2 | *STAT2* | N | Eurasians, Oceanians | Immune system regulation | Mendez et al. 2012 |
| 1:117-122MB | TBX15 | *TBX15/WAR2* | D | Native Americans | Adipose tissue differentiation | Racimo et al. 2017 |
| 4:37-42MB | TLR1610 | *TLR1610* | N | Europeans | Immune system regulation | Gittelman et al. 2016, Descamps et al. 2016 |
| 6:135.5-140.5MB | TNFA1P3 | *TNFA1P3* | N, D | Oceanians | Immune system regulation | Gittelman et al. 2016 |
| 9:110-115MB | TXN | *TXN* | N | East Asians | Redox reactions | Browning et al. 2018 |

*This tables shows the adaptive introgression candidate regions identified in modern humans that are used in this study. The first column indicate the 5MB region range with format of chromosome:coordinate range (Hg19). The second column shows the identifier of the region in this study, and the third column shows the AI candidate genes included in the region. The fourth column shows the identified archaic introgression group, with N = Neanderthals and D = Denisovans. The fifth column shows the modern human populations where the AI signature are identified. The sixth column shows the biological pathways involved by the genes. The original studies reported the AI signature are shown in the last column.*

**Supplementary Figure 1: Exon density distribution in human genome (5MB windows)**

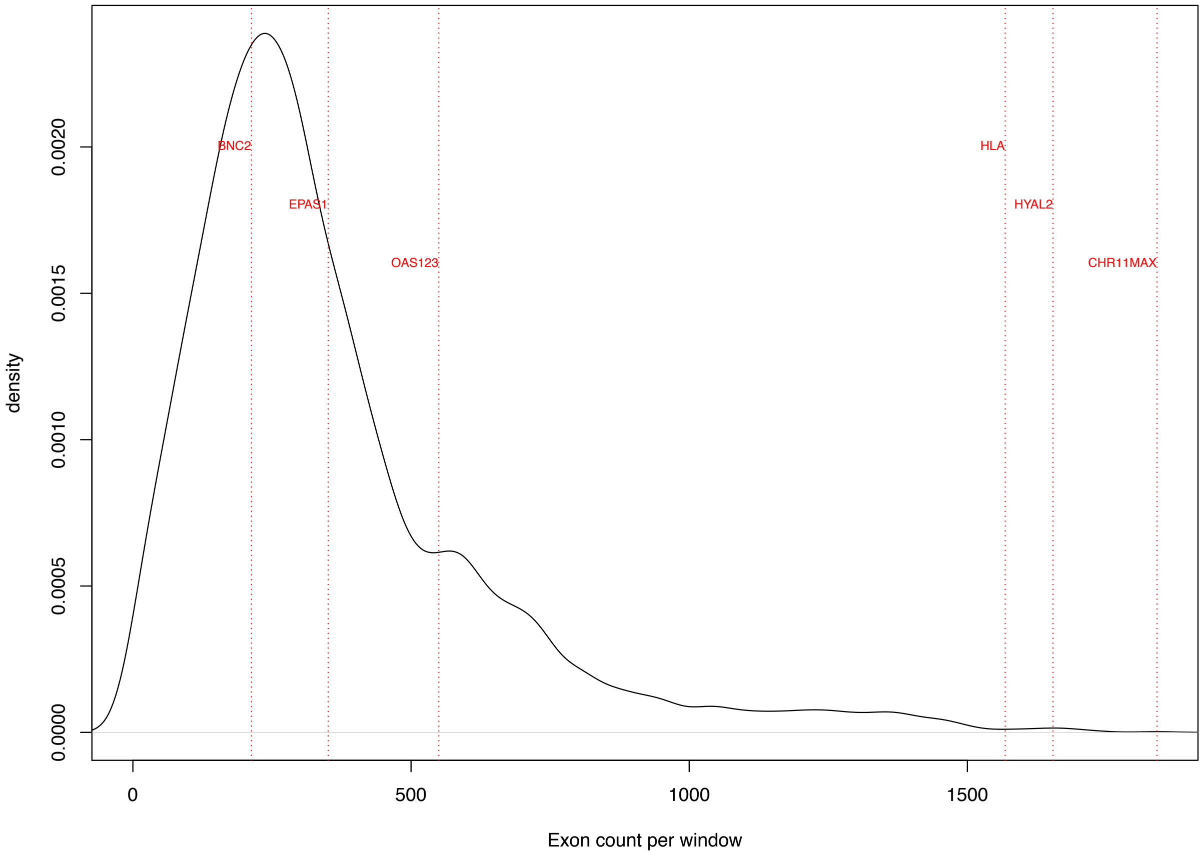


*This figure shows the exon density distribution of 5MB-windows in human genome. The exon density is defined as the number of exons within a 5MB window. The whole human genome was scanned in 5MB sliding windows with 1kb step size, and the exon density distribution is plotted as the black solid line. Each vertical red dotted line shows the exon density of the AI candidiate regions that are mentioned in the main text (identifiers shown in Supp. Table 1).*

**Supplementary Figure 2: AI signature statistics distribution under Model_0 using uniform low recombination rates at 1e-9 (*Chr11max* region)**

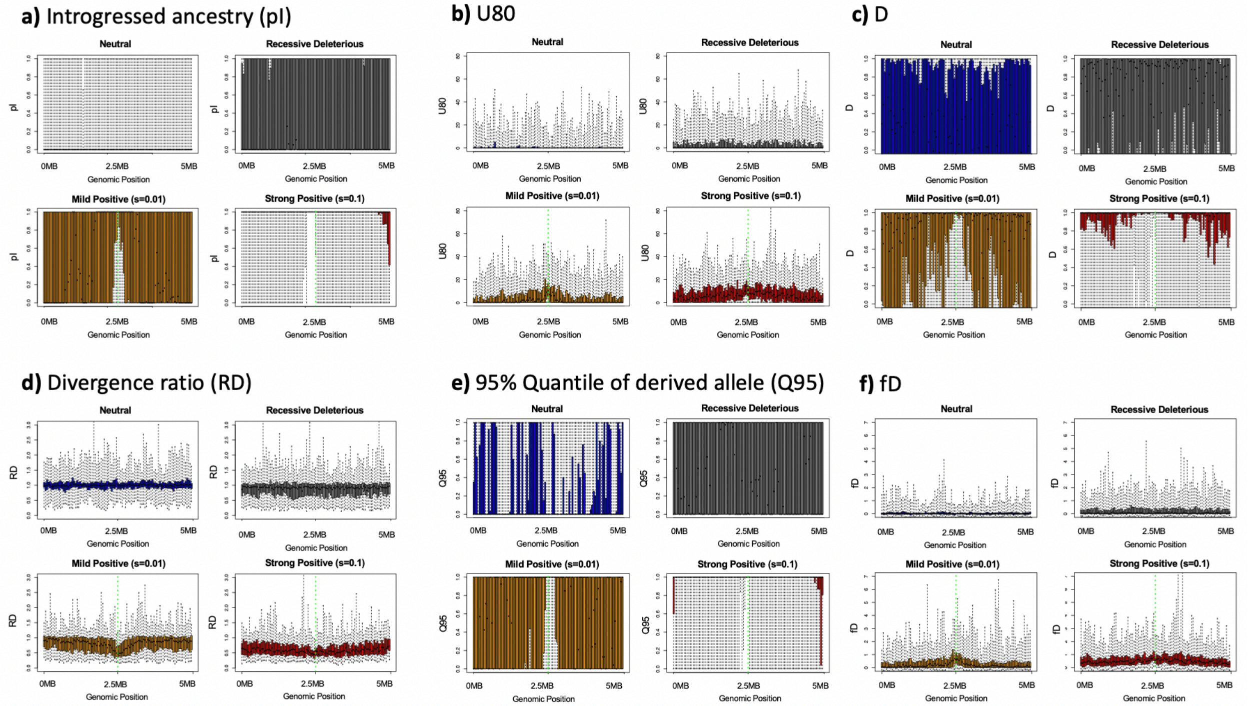


*From panel a-f, the distribution of AI statistics from Chr11max region (demographic model = Model_0) simulation from the four mutation models are shown in order of pI, U80, D, RD, Q95, and fD (definition of statistics see main text Table 1). The neutral, deleterious, Mild-Pos, and Strong-Pos models are illustrated in blue, gray, orange and red respectively. Recombination was simulated at a uniform rate of 1e-9. The adaptive mutations in the latter two mutation models are introduced in window in the middle of the region (2.5MB), indicated by the green solid line. The plot shows the interquartile distributions of U80 in boxes, with whiskers extend to all data points.*

**Supplementary Figure 3: False positive rate distribution of AI signature statistics under Model_0 using uniform low recombination rates at 1e-9 (*Chr11max* region)**

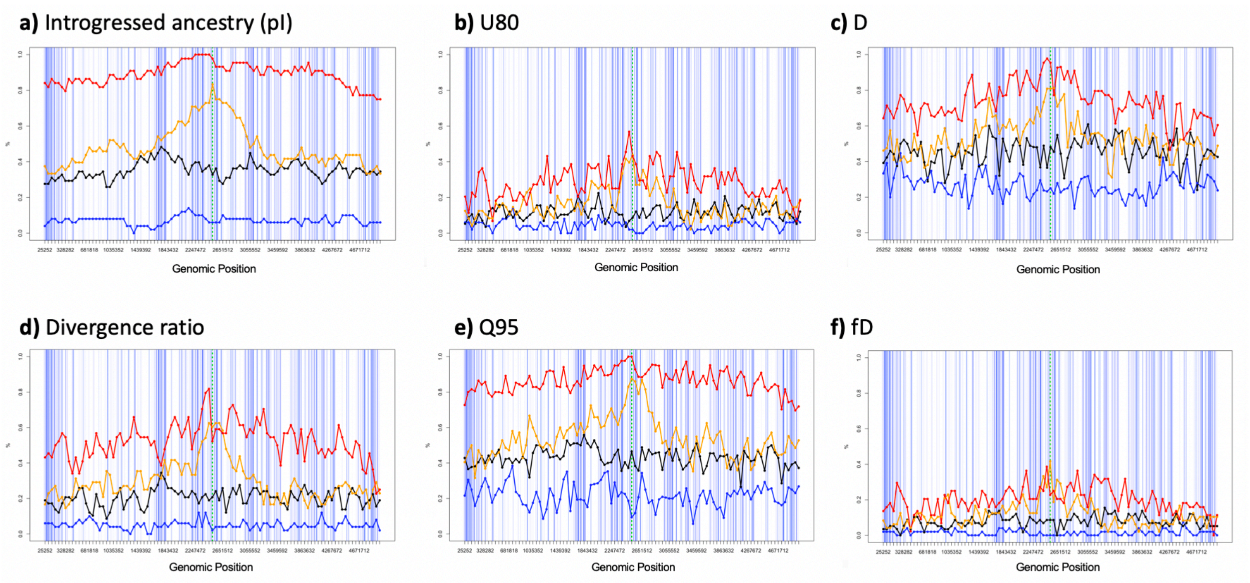


*This figure the False Positive Rates and True Positive Rates for summaries of the data in Supp. Fig. 2. The positive rates are computed against the significance threshold value defined as the most-extreme 5% value from Neutral distribution, and we calculate FPR or TPR using the percentage of statistical values per 50kb-window that exceeds the threshold value. For the simulations, red, orange, blue and black represent Strong-Pos, Mild-Pos, Neutral, and Deleterious respectively. The light blue lines in the right-panels illustrate the exons where new mutations can arise, and the green solid line represents the window where the adaptive mutation occurred. The simulations ran under Model_0 using the genic structure of the Chr11Max region, using a uniform low recombination rate of 1e-9.*

**Supplementary Figure 4: False positive rate of pI, RD, and U80 under Model_0 using realistic recombination rate (*Chr11max* region)**

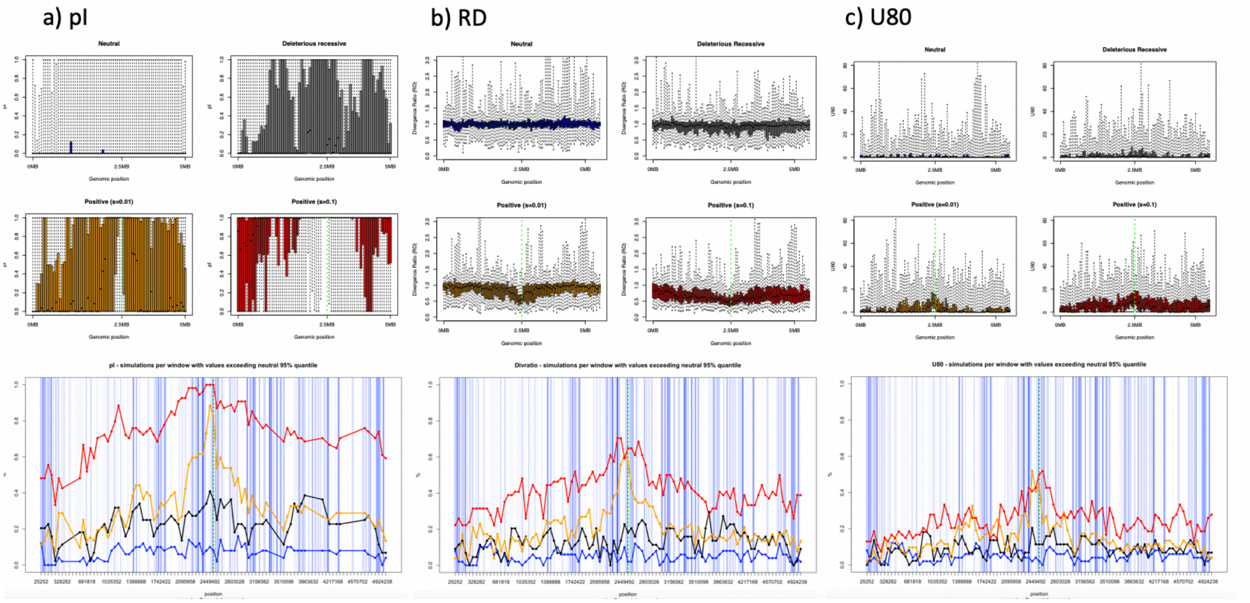


*This figure captures the distribution and False/True positive rates in three statistics: pI, RD, and U80. The simulations are performed using the Chr11max region under Model_0, with the realistic recombination map for modern humans (in contrast to Supp. Fig. 2-3). The color codes correspond to the definition in Supp Fig. 2-3.*

**Supplementary Figure 5: False positive rate of pI, RD, and U80 under Model_h using realistic recombination rate (*Chr11max* region)**

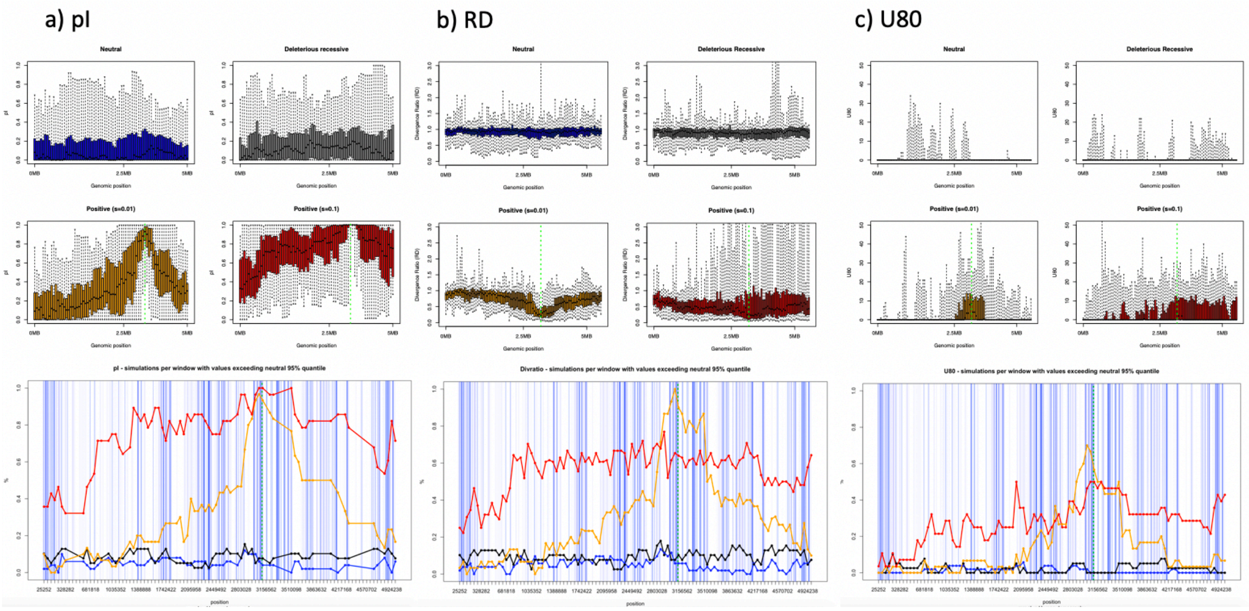


*This figure captures the distribution and False/True positive rates in three statistics: pI, RD, and U80. The simulations are performed using the Chr11max region under an approximate human demographic model (Model_h), with the realistic recombination map for modern humans. The color codes correspond to the definition in Supp Fig. 2-3.*

**Supplementary Figure 6: False Positive Rates on signature statistics from human AI candidate regions under model_h using realistic recombination rates**

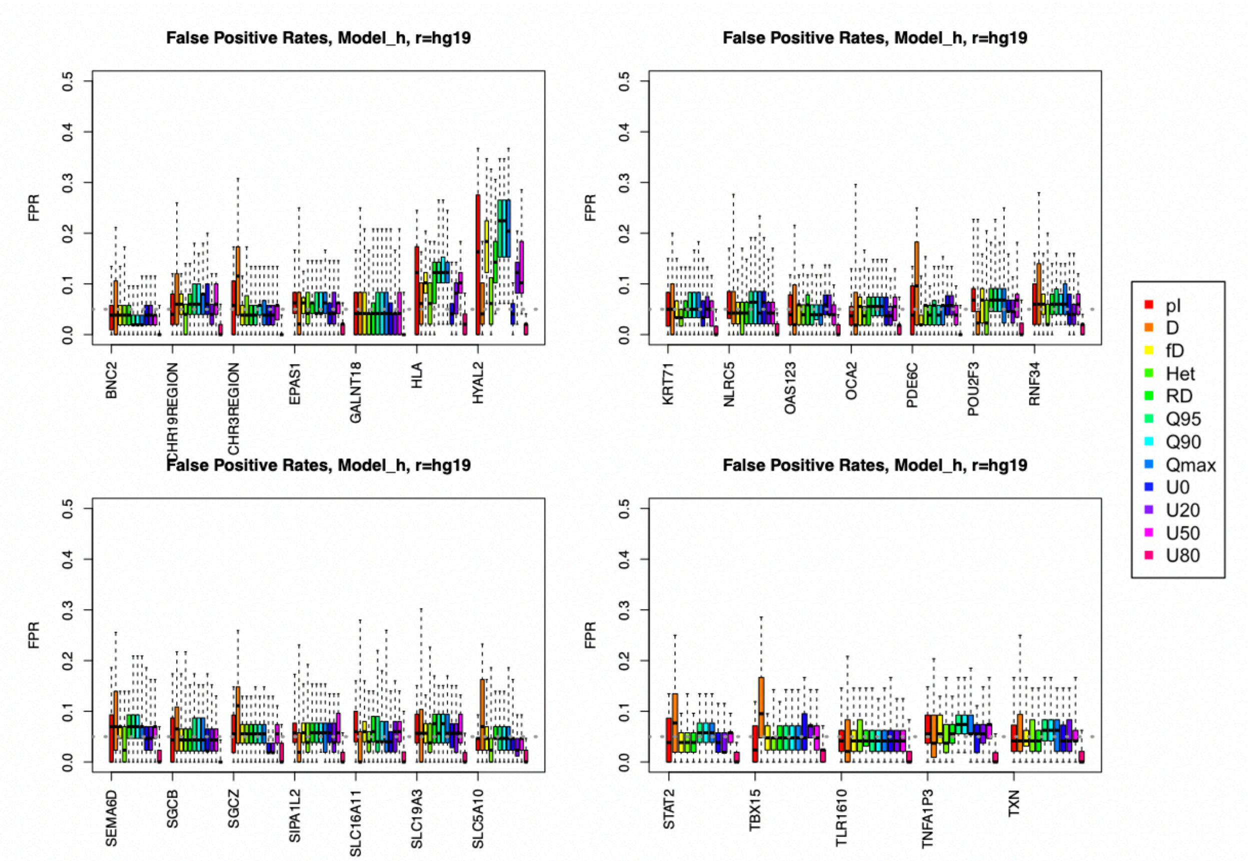


*The summary statistics are obtained from simulations under Neutral and Deleterious mutation models using human demography, Model_h. The recombination rates and exon density reflect the four regions in human genome harboring the 26 AI candidate genes. The FPR (y-axis) is computed assuming a neutral null model and represents the proportion of simulations replicates under the Deleterious model that are called significant for adaptive introgression.*

**Supplementary Figure 7. Demography illustration of four different recipient population growth patterns in recent human history**

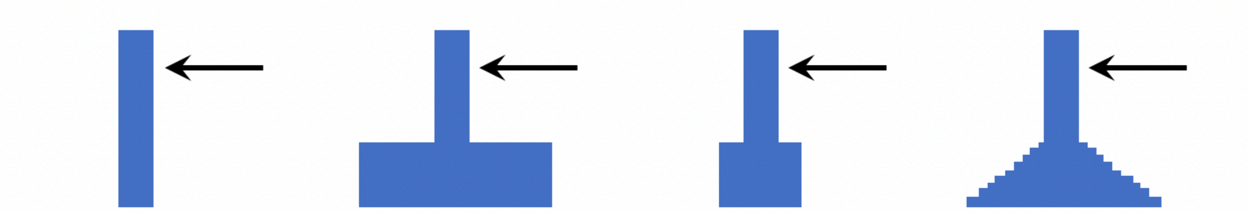


*Four different demography scenarios were used to test if the recent growth in human history (Eurasian population) played a role in contributing to the false positive rates in AI detection due to the heterosis effect. Specifically, the "growth 4" is the same as in Model_h (main text Fig. 2b) where the recipient population ("pR" or "Eurasian") experiences an exponential growth that the population size at the end of the simulation ($N_{end}$) is 41080. As comparison, "Growth 1" experiences no population growth and remains the size since the bottleneck; "Growth 2" experiences an immediate population growth where the population size matches with the end population size in "Growth 1"; "Growth 3" is an intermediate situation between Growth 1 and 2 that the recipient population experiences an inmmediate growth while the size change is limited. The rest of the demography remains the same as in Model_h, and the AI candidate regions are simulated under each growth pattern scenario for 200 replicates.*

**Supplementary Figure 8. False population rates of genes in main text Fig. 6 at Growth patterns 1-4**

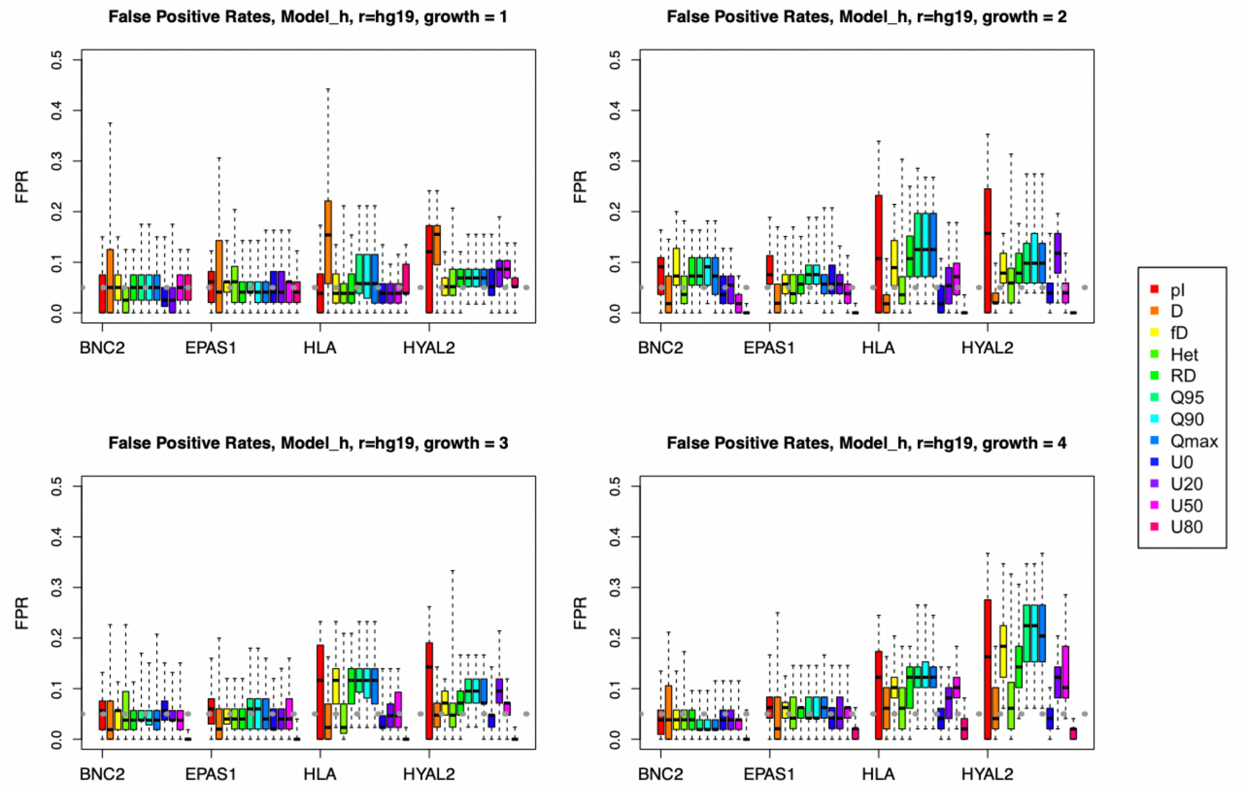


*The "typical genes" (BNC2 and EPAS1) and the "outlier genes" (HYAL2 and HLA) mentioned in main text Fig 6 are simulated under the 4 growth patterns at the "neutral" and "deleterious" mutation scenarios each for 200 replicates. The rest of the demography is the same as Model_h, and the realistic recombination rate map was applied in the simulations. Here it shows that the outlier genes remain high false positive rates despite the changes in recipient population growth pattern. And the situations where the recipient population size at the end of the simulation is large (Growth 2 and 4) result in slightly higher FPRs especially in the outlier genes than the situations where recipient population size remain small (Growth 1 and 3).*

**Supplementary Figure 9: False positive rates of genes in main text Fig. 6 with different recombination rates**

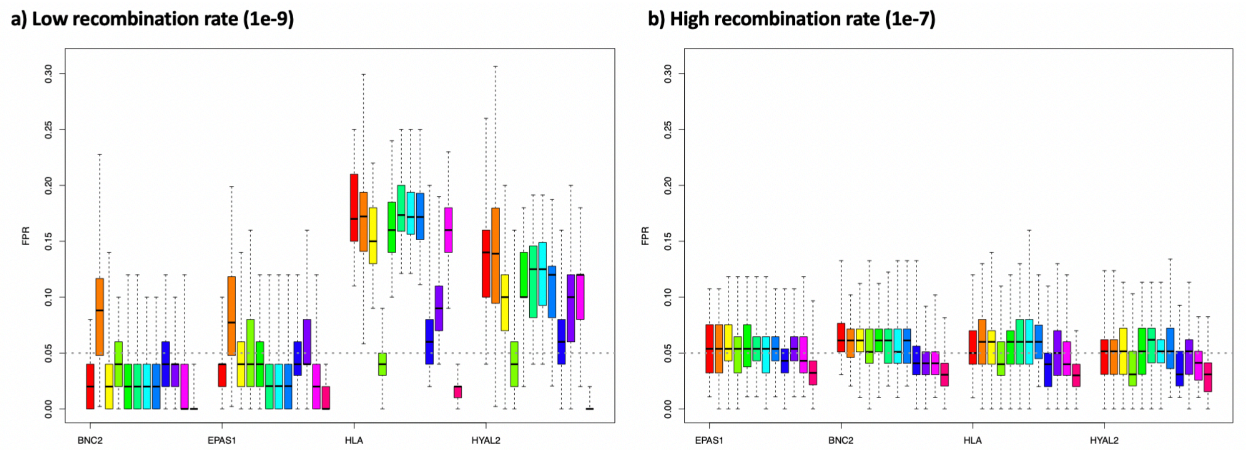


*The previously mentioned 4 genes from main text Fig. 6 are simulated under the "neutral" and "deleterious" mutation models with two different uniform recombination rates (low rate at 1e-9, left panel; and high rate at 1e-7, right panel) each for 200 replicates under Model_h. Here it shows that a uniformly low recombination rate does not increase the FPRs in "typical genes" on most of the statistics with an exception of D statistics. The low recombination rate also amplifies the FPRs in the outlier genes further than previously seen. And when a uniformly high recombination rate is applied, the FPRs resume low for all genes. The color codes for the summary statistics are the same as Supp. Fig 7-8.*

**Supplementary Figure 10:** Exon density and mean recombination rate distribution in human genome

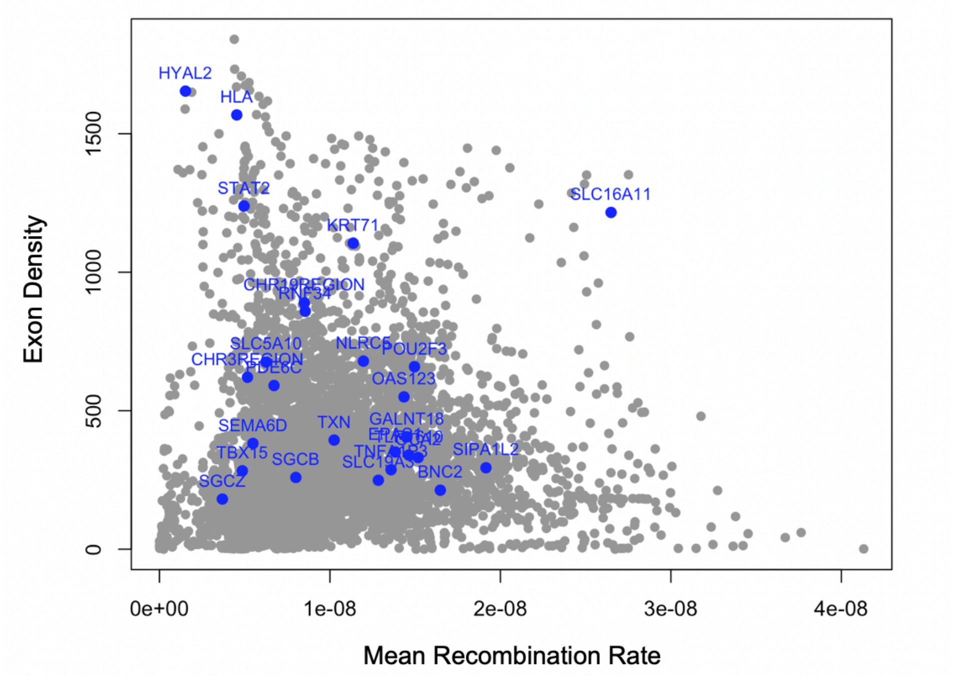


*This figure shows the relationship between the exon density and mean recombination rate in sliding 5MB windows (gray points) across the human genome (window definition see Supp. Fig. 1). The blue points highlight all the AI candidate regions mentioned in Supp. Table 1.*

**Supplementary Figure 11. P-values between neutral and deleterious null models on AI candidate regions**

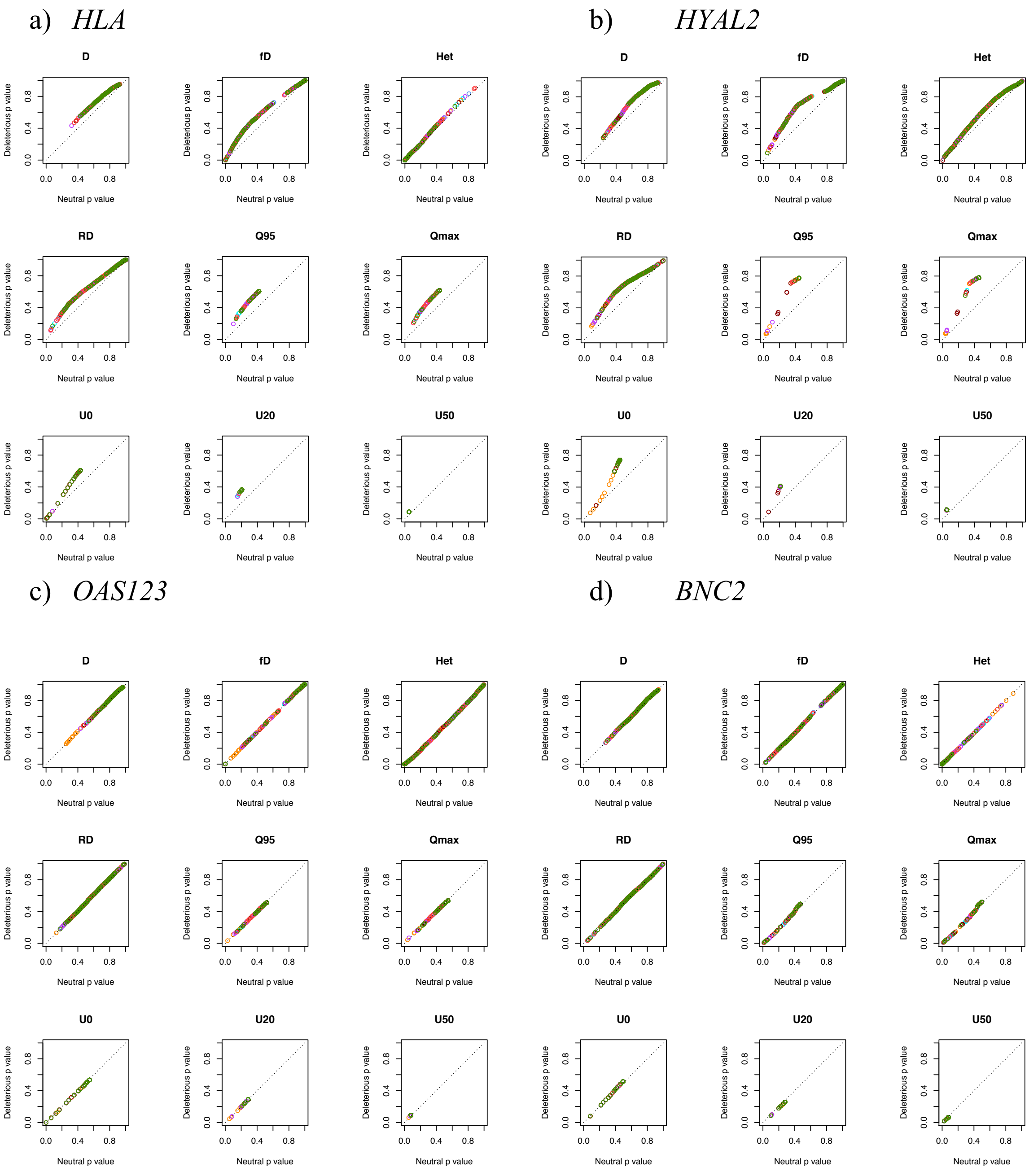


*For each 50kb-window in the empirical data, we calculated the p-value using neutral null distribution (x-axis) and deleterious null distribution (y-axis) respectively, and plotted the relationship between the two p-values as a single point. Different colors represent the 7 populations from 1000 genomes data. The points above the diagonal indicate the windows where deleterious null models made more conservative predictions in corresponding populations. Color codes for 1000 Genomes populations see main text Fig. 8 and Supp. Fig. 12.*

#### Supplementary Figure 12: Significance window number difference in AI candidate regions between different null models

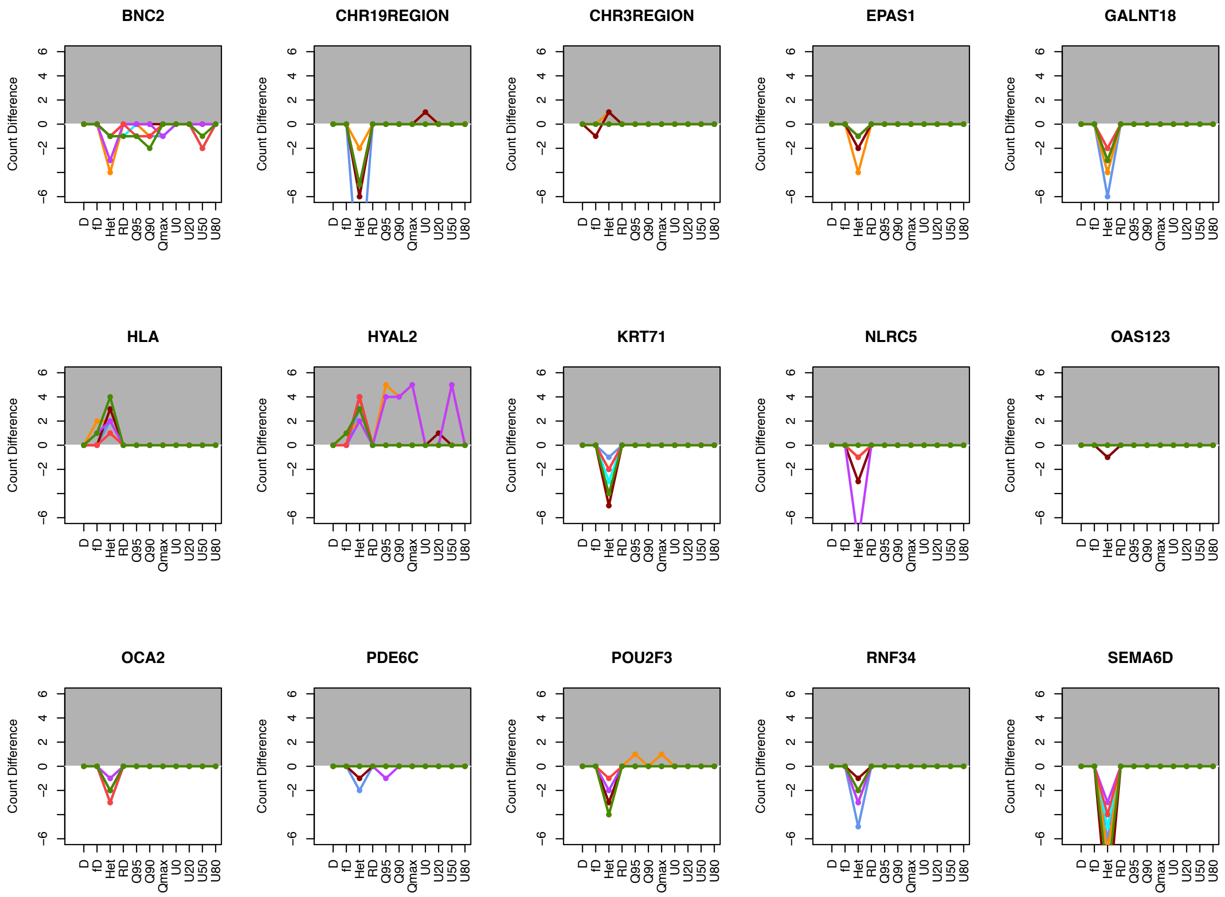

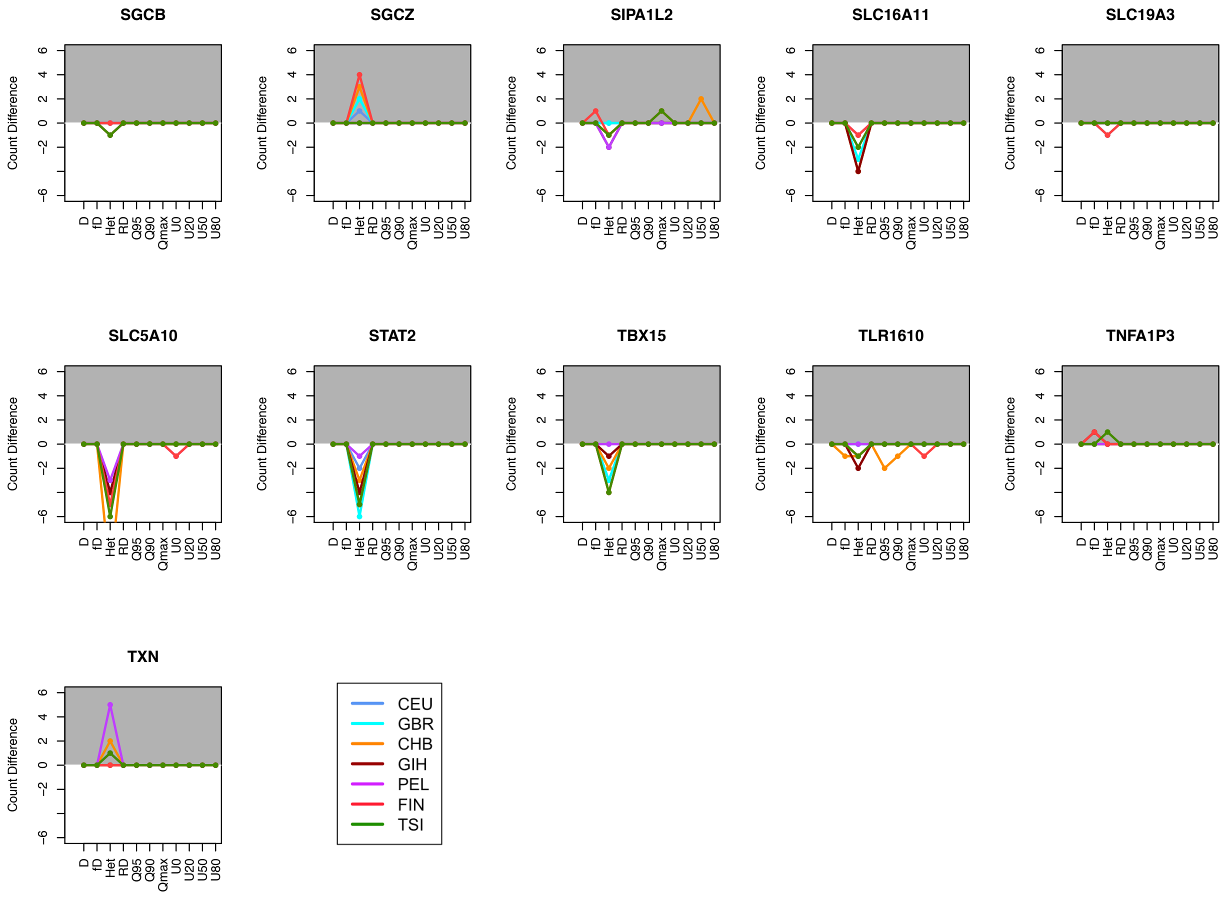


*We compared the difference in the number of significant hits (windows with p-value <0.05) for all 26 AI candidate regions predicted by neutral and deleterious null models. Each point represents that difference in number (y-axis; Neutral significance number – Deleterious significance number) in its corresponding statistics (x-axis). The genes with multiple points above y-axis value 0 are highlighted in the gray-shaded area, indicating the deleterious null models predict fewer window hits being significant for given statistics, which implies potential false positives from neutral null models.*

**Supplementary Figure 13: Significance window numbers in *HLA* and *HYAL2* genes under neutral and deleterious null models**

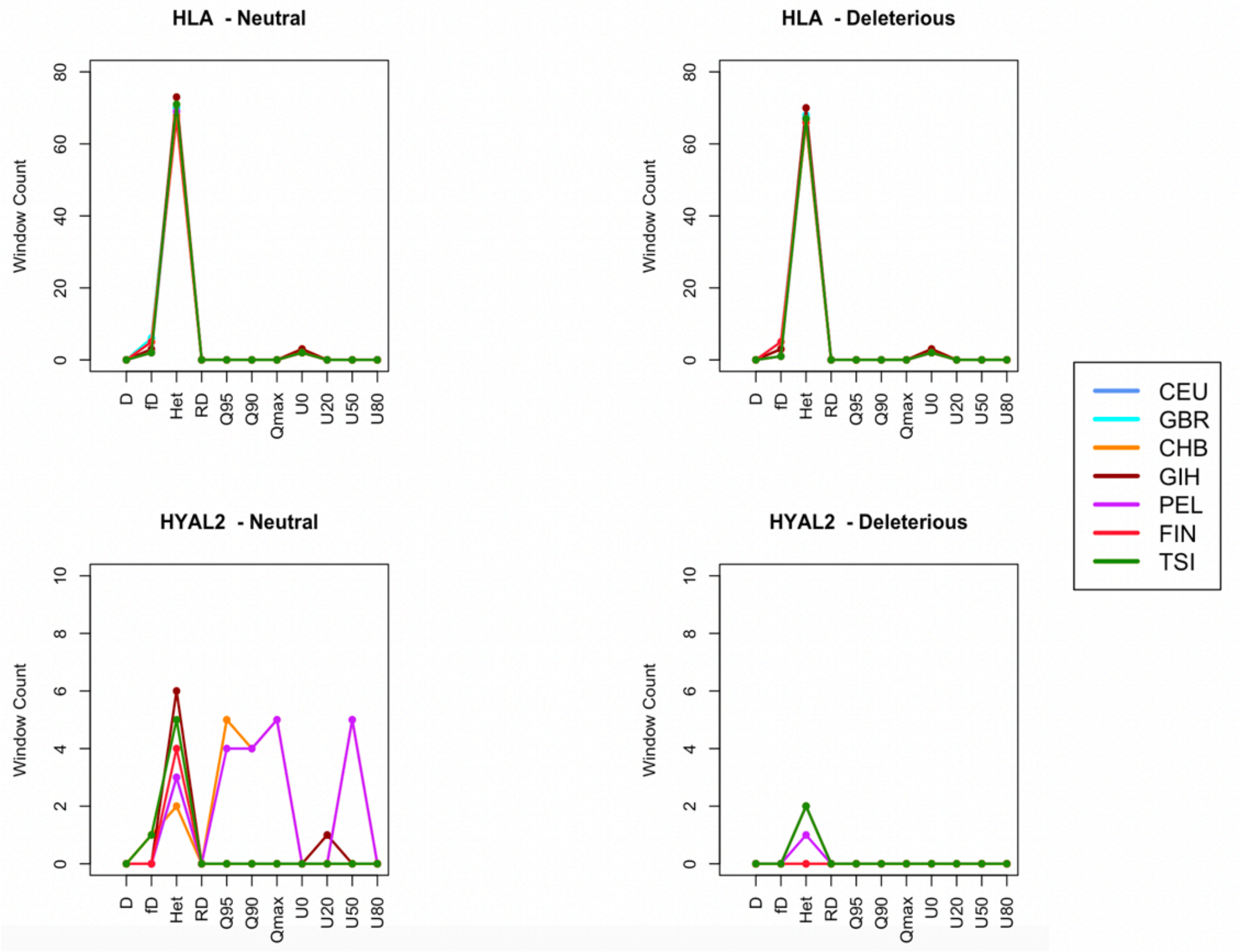


*This figure shows the number of significant hits (windows with p-value <0.05) for the HLA and HYAL2 gene regions predicted by neutral (left panels) and deleterious (right panels) null models. Each point represents that number of significance (y-axis) in its corresponding statistics (x-axis).*
